# Afferent projections to the *Calca*/CGRP-expressing parabrachial neurons in mice

**DOI:** 10.1101/2024.05.07.593004

**Authors:** Mustafa Korkutata, Roberto De Luca, Bridget Fitzgerald, Elda Arrigoni, Thomas E. Scammell

## Abstract

The parabrachial nucleus (PB), located in the dorsolateral pons, contains primarily glutamatergic neurons which regulate responses to a variety of interoceptive and cutaneous sensory signals. The lateral PB subpopulation expressing the *Calca* gene which produces the neuropeptide calcitonin gene-related peptide (CGRP) relays signals related to threatening stimuli such as hypercarbia, pain, and nausea, yet the afferents to these neurons are only partially understood. We mapped the afferent projections to the lateral part of the PB in mice using conventional cholera toxin B subunit (CTb) retrograde tracing, and then used conditional rabies virus retrograde tracing to map monosynaptic inputs specifically targeting the PB^*Calca*/CGRP^ neurons. Using vesicular GABA (vGAT) and glutamate (vGLUT2) transporter reporter mice, we found that lateral PB neurons receive GABAergic afferents from regions such as the lateral part of the central nucleus of the amygdala, lateral dorsal subnucleus of the bed nucleus of the stria terminalis, substantia innominata, and the ventrolateral periaqueductal gray. Additionally, they receive glutamatergic afferents from the infralimbic and insular cortex, paraventricular nucleus, parasubthalamic nucleus, trigeminal complex, medullary reticular nucleus, and nucleus of the solitary tract. Using anterograde tracing and confocal microscopy, we then identified close axonal appositions between these afferents and PB^*Calca*/CGRP^ neurons. Finally, we used channelrhodopsin-assisted circuit mapping to test whether some of these inputs directly synapse upon the PB^*Calca*/CGRP^ neurons. These findings provide a comprehensive neuroanatomical framework for understanding the afferent projections regulating the PB^*Calca*/CGRP^ neurons.

## 1 INTRODUCTION

The parabrachial nucleus (PB) coordinates responses to a variety of interoceptive and cutaneous signals. Within the external lateral subnucleus of the PB (PBel), neurons expressing *Calca* which encodes for calcitonin gene-related peptide (CGRP) are especially important for responding to a variety of threatening stimuli, including pain, nausea, and hypercarbia, and then driving appropriate behavioral responses via projections to the forebrain and brainstem. For example, the PB^*Calca*/CGRP^ neurons receive pain signals from the dorsal horn, leading to cortical activation and waking in response to pain (Saper, 2016; Kaur et al., 2017; Palmiter, 2018; Lynch et al., 2022; Alexandre et al., 2024). They are also necessary for triggering arousals in response to hypercapnia via projections to the amygdala, basal forebrain and lateral hypothalamus (Kaur et al., 2017). In addition, the PB^*Calca*/CGRP^ neurons have been implicated in various fear responses and the formation of aversive memories, plus satiation and conditioned taste aversion (Carter et al., 2013, 2015; Campos et al., 2016, 2017, 2018; Kaur et al., 2017; Chen et al., 2018; Palmiter, 2018). More broadly, the PB^*Calca*/CGRP^ neurons may provide a general alarm signal (Saper, 2016; Palmiter, 2018), so defining what regulates these neurons is essential for developing better therapies for managing pain, nausea, hypoventilation, and other conditions.

Prior work using conventional anterograde and retrograde tracing has shown that the PB^*Calca*/CGRP^ neurons and the external lateral PB receive inputs from a variety of CNS regions, including the spinal cord, nucleus of the solitary tract (NTS), reticular formation, periaqueductal gray (PAG), superior and inferior colliculi, arcuate nucleus (Arc), lateral dorsal bed nucleus of the stria terminalis (BSTLD), and central nucleus of the amygdala (CeA) (Palmiter, 2018; Kang et al., 2022; Kirouac et al., 2022) (**Table 4** provides a list of all abbreviations which follow Paxinos (Paxinos and Franklin, 2004) except for some abbreviations from Swanson (Swanson, 2004)). The PB^*Calca*/CGRP^ neurons receive inhibitory inputs from agouti-related protein (AgRP) neurons of the arcuate nucleus (Arc) (Campos et al., 2016), but the mapping and characterization of other inputs remain incomplete. In fact, most prior work used conventional tracing, so it remains unclear just which inputs directly regulate the activity of the PB^*Calca*/CGRP^ neurons.

To address this gap, we used conventional and conditional retrograde and anterograde tracers to systematically map inputs to the *Calca*/CGRP neurons in the lateral PB of mice. We then used GABA (vGAT) and glutamate (vGLUT2) reporter mice to define the neurochemical identity of these inputs. In addition, we used channelrhodopsin-2 (ChR2)-assisted circuit mapping (CRACM) to test whether some inputs monosynaptically innervate the PBel^*Calca*/CGRP^ neurons.

## 2 MATERIALS AND METHODS

### 2.1 Animals

These experiments were approved by the Institutional Animal Care and Use Committee of Beth Israel Deaconess Medical Center and were performed in accordance with the National Institutes of Health Guide for the Care and Use of Laboratory Animals.

We used *C57BL/6J* mice (JAX, 12-24 weeks old) (**Table 1**) to map inputs to the lateral PB area using the conventional retrograde tracer cholera toxin B subunit (CTb). To identify monosynaptic inputs to the PB^*Calca*/CGRP^ neurons using conditional rabies virus tracing, we used *Calca-Cre* mice (*Calca*^Cre:EGFP^;B6.CgCalcatm1.1(Cre/EGFP)Rpa/J, JAX Cat: #033168) with Cre:EGFP inserted into Exon 2 of the endogenous *Calca* gene (Carter et al., 2013). To determine the neurochemical identity of neurons projecting to the lateral PB, we used vesicular GABA transporter (vGAT) reporter mice (*vGAT-IRES-Cre::R26-lox-STOP-lox-L10-GFP*). The vGAT and vGLUT2 reporter mice were generated by crossing *vGAT-IRES-Cre* mice and *vGLUT2-IRES-Cre* mice with *R26-lox-STOP-lox-L10-GFP* mice, respectively. For anterograde labeling of inputs to the lateral PB with cre-dependent expression of ChR2, we used *vGAT-IRES-Cre* or *vGLUT2-IRES-Cre mice*. For functional testing of inputs to PBel^*Calca*/CGRP^ neurons with CRACM, we used *Calca-Cre::vGAT-Flp* mice. The *Calca-Cre::vGAT-Flp* mice express Cre-recombinase in *Calca*-CGRP-expressing neurons and Flp-recombinase in vGAT-expressing neurons and were produced by crossing the *Calca*^Cre:EGFP^ (JAX, Cat: #033168) with the Slc32a1tm1.1(flpo)Hze mice (vGAT-Flp; B6.Cg-Slc32a1tm1.1(flpo)Hze/J, JAX Cat. #029591). All mice were backcrossed to *C57BL/6J* for at least 5 generations. Mice were 12-24 weeks old at the time of experiments, and littermates of either sex were assigned to experimental groups. After surgery, mice were individually housed in standard plastic cages with corncob bedding with nesting materials on a 12h light:12h dark cycle at ambient temperatures ranging of 22 ± 2°C. Mouse chow (Teklad F6 Rodent Diet 8664) and water were provided *ad libitum*.

**TABLE 1.** Mouse strains used in this study.

| Mouse strains | Source | References |
| --- | --- | --- |
| C57BL/6J | JAX 000664 | Simon et al., 2013 |
| Calca-Cre<br>(Calca<sup>Cre:GFP</sup>) | JAX 033168 | Carter et al., 2013 |
| vGAT-IRES-Cre | JAX 016962 | Vong et al., 2011 |
| vGAT-IRES-Flp | JAX 02591 | Jo et al., 2018 |
| vGLUT2-IRES-Cre | JAX 016963 | Vong et al., 2011 |
| L10-GFP or R26-lox-STOP-lox-L10-GFP | B. Lowell, BIDMC, Harvard University | Krashes et al., 2014 |

### 2.2 Stereotaxic Injections

We anesthetized mice with ketamine/xylazine (Patterson Vet, Product # 07-803-6637 and #07-808-1947, 100/10 mg/kg, i.p.) and placed them in a stereotaxic alignment system (Model 1900; Kopf Instruments). We made a midline incision, identified bregma, and drilled a burr hole. We then injected tracers or AAV via a glass micropipette (∼20 μm tip diameter) using an air pressure microinjection system. The glass micropipette was left in place at the injection site for an additional 5-7 min, then slowly withdrawn before closing the incision with Vetbond (3M). For postoperative analgesia, we administered meloxicam (Zoo Pharm, Product # 1Z-88000, 4mg/kg, s.c.).

### 2.3 Retrograde tracing with CTb

To retrogradely label inputs to the lateral PB area, we unilaterally injected the lateral PB (Anterior-Posterior (AP): -5.3 mm, Medial-Lateral (ML): -1.4 mm, Dorsal-Ventral (DV): 3.4 mm) of *C57BL/6J* mice (n=4) with 3 nL of CTb (List Biological Labs, Cat#103A, Campbell, CA; 1% in saline). To determine the neurochemical identity of lateral PB-projecting neurons, we similarly microinjected 3 nL CTb into the lateral PB of *vGAT-* and *vGLUT2-reporter* mice (n=4, each). We perfused mice one week later for histology.

### 2.4 Retrograde tracing with conditional rabies virus

To map monosynaptic inputs to the PB^*Calca*/CGRP^ neurons, we used conditional rabies virus tracing, which restricts rabies virus infection and replication through Cre-dependent expression of the TVA gene (avian leukosis and sarcoma virus subgroup A receptor, chicken) and rabies glycoprotein G (RG). We unilaterally injected the lateral PB of *Calca-Cre* mice (n=11) with 75 nL of *AAV8-Ef1a-FLEX-TVA-mCherry* (UNC Vector Core, Lot#AV5008b, Titer: 5.4x10^12^ μg/mL) and *AAV2/8-CAG-FLEX-RG-hGH* (Stanford Gene Vector and Virus Core, Lot#AAV59, Titer: 2.09x10^13^ μg/mL) in a 1:1 ratio (**Table 2**). Four weeks later, we injected the lateral PB of these mice with 90 nL of conditional rabies virus (EnvA SADΔG-EGFP; Salk Institute, Addgene#32635, Titer: 2.26x10^8^ TU/mL). We then perfused the mice 7 days later.

**TABLE 2.** Virus strains and recombinant protein used in this study.

| Virus strains/<br>Recombinant<br>Protein | Source | Titer |
| --- | --- | --- |
| AAV2/8-CAG-FLEX-RG-hGH | Stanford,<br>Lot#AAV59 | $2.09 \times 10^{13}$<br>μg/mL |
| AAV8-Ef1a-DIO-hChR2 (H134R)-mCherry | UNC,<br>Lot#AV9080 | $1.4 \times 10^{13}$<br>μg/mL |
| AAV8-Ef1a-FLEX-TVA-mCherry | UNC,<br>Lot#AV5008b | $5.4 \times 10^{12}$<br>μg/mL |
| AAV8-hSyn-DIO-mCherry | Bryan Roth,<br>Addgene,<br>Catalog#50459 | $2.2 \times 10^{13}$<br>GC/ml |
| AAV9-CAG-hChR2-mCherry | Karl Deisseroth,<br>Addgene,<br>Catalog#100054 | $3.3 \times 10^{13}$<br>GC/ml |
| AAVd/j-Ef1a-fDIO-hChR2-EGFP | UNC | $2 \times 10^{12}$<br>virus<br>mol./ml |
| Cholera Toxin<br>Subunit B | List Biological<br>Labs, Cat#103A<br>Campbell, CA | 1% in<br>saline |
| SADΔG-EGFP | Salk Institute,<br>Addgene,<br>Catalog#32635 | $2.26 \times 10^8$<br>TU/mL |

### 2.5 Conditional anterograde tracing

To determine which brain regions project to the PBel^*Calca*/CGRP^ neurons, we performed conditional anterograde tracing using microinjections of 6-60 nL of *AAV8-Ef1a-DIO-hChR2 (H134R)-mCherry* (UNC, Lot# AV9080, Titer: 1.4x10^13^ μg/mL). We unilaterally injected this AAV into the CeA (AP: -1.5 mm, ML: -2.8 mm, DV: 4.7 mm, n=4) (Paxinos and Franklin, 2004), bed nucleus of the stria terminalis (BST) (AP: -1.2 mm, ML: -0.2 mm, DV: 4.0 mm, n=4), substantia innominata (SI) (AP: -0.1 mm, ML: -1.8 mm, DV: 4.8 mm, n=4), and ventrolateral periaqueductal gray (VLPAG) (AP: - 4.6 mm, ML: -0.5 mm, DV: 2.6 mm, n=4) of *vGAT-IRES-Cre* mice.

We also injected this AAV into the paraventricular nucleus of the hypothalamus (PVH) (AP: -0.8 mm, ML: -0.2 mm, DV: 4.6 mm, n=4), parasubthalamic nucleus of the hypothalamus (PSTh) (AP: -2.3 mm, ML: -1.2 mm, DV: 4.6 mm, n=4), and NTS (AP: -6.2 mm, ML: - 1.2 mm, DV: 2.7 mm, n=4) of *vGLUT2-IRES-Cre* mice.

To trace inputs from the infralimbic cortex (IL) (AP: 1.9 mm, ML: -0.6 mm, DV: 2.9 mm, n=4) and insular cortex (IC) (AP: -0.6 mm, ML: -3.3, DV: 3.3 mm, n=4), we first used a conditional tracing approach by injecting 3-9 nL of *rAAV5-CAMKIIα-hChR2-mCherry* in *C57BL/6J* mice, but as this failed to label axons in the PB, we used a non-conditional tracing approach by injecting 100 nL of *AAV9-CAG-hChR2-mCherry* (Addgene, Catalog#100054-AAV9, Titer: 3.3x10^13^ GC/ml) in *C57BL/6J* mice. We perfused mice 3-4 weeks later to allow sufficient time for ChR2-mCherry expression and anterograde tracing. Using confocal microscope with 1 μm optical sections, we mapped axon terminals closely apposing the soma and dendrites of PB^*Calca*/CGRP^ neurons.

### 2.6 In vitro electrophysiology

We used ChR2-assisted circuit mapping (CRACM) to determine whether the CeA GABA neurons directly inhibit the PBel^*Calca*/CGRP^ neurons (CeA^GABA^→ PBel^*Calca*/CGRP^). Using *vGAT-Flp::Calca-Cre* mice (n = 5), we unilaterally injected the CeA (AP: -1.5 mm, ML: -2.8, DV: 4.7 mm) with 80 nl of *AAVDJ-Ef1a-fDIO-ChR2-eYFP* (UNC Gene Therapy Center, Titer: 2 × 10^12^ virus molecules/ml) and placed a second microinjection of 200 nl of *AAV8-hSyn-DIO-mCherry* (Addgene, Catalog#50459-AAV8, Titer: 2.2x10^13^ GC/ml) into the ipsilateral PB to label the PB^*Calca*/CGRP^ neurons (AP: -5.3 mm, ML: -1.4 mm, DV: 3.4 mm). Four to six weeks after the AAV injections, we prepared coronal brain slices (250 μm-thick) containing the PB and recorded from mCherry-labeled PBel^*Calca*/CGRP^ neurons using a combination of fluorescence and infrared differential interference contrast (IR-DIC) microscopy. Using a fixed stage upright microscope (BX51WI, Olympus America Inc) equipped with a Nomarski water immersion lens (Olympus 40X / 0.8 NAW) and IR-sensitive CCD camera (ORCA-ER, Hamamatsu, Bridgewater, NJ), we acquired real time images using Micro-Manager software.

We recorded mCherry-labeled PBel^*Calca*/CGRP^ neurons in brain slices that were maintained submerged and perfused (1.5 ml/min) with artificial cerebrospinal fluid (ACSF). Recordings were conducted in whole-cell configuration using a Multiclamp 700B amplifier (Molecular Devices, Foster City, CA), a Digidata 1322A interface, and Clampex 9.0 software (Molecular Devices). Neurons were excluded from analysis if their input resistance changed more than 10% over the recording. We photostimulated ChR2-expressing CeA^GABA^ terminals and axons in the PB using full-field light (∼10 mW/mm2, 1 mm beam width) from a 5 W LUXEON blue light-emitting diode (470 nm wavelength; #M470L2-C4; Thorlabs, Newton, NJ) coupled to the epifluorescence pathway of the microscope. We photo-evoked inhibitory postsynaptic currents (IPSCs) with 10 ms light pulses (0.1 Hz, for a minimum of 30 trials). We recorded photo-evoked IPSCs in ACSF containing 1 mM of kynurenic acid, in whole-cell voltage-clamp mode at a holding potential of 0 mV using a Cs-methane-sulfonate-based pipette solution. The pipette solution also contained 0.5% biocytin to label the recorded neurons. We purchased kynurenic acid from Cayman Chemical (Ann Arbor, MI), and all other chemicals from Fisher Scientific (Waltham, MA) or Sigma-Aldrich (Saint Luis, MO). We analyzed recordings using Clampfit 10 (Molecular Devices) and Python 3 (www.python.org) and software. To ensure unbiased detection of the synaptic events, the IPSCs were detected and analyzed automatically using MiniAnalysis 6 software (Synaptosoft, Leonia, NJ). We considered PBel^*Calca*/CGRP^ neurons to be responsive to photostimulation if their IPSC probability during the first 50 ms after the light pulses was greater than baseline IPSC probability plus five times the SEM (baseline IPSC probability = 7.25 ± 0.12, *n* = 26). We calculated the latency of the photo-evoked IPSCs as the time difference between the start of the light pulse and the 5% rise point of the first IPSC (Hull et al., 2009). We present data as mean ± SEM, and *n* refers to the number of cells per group. Group means were compared using *paired* t-tests. Values indicating p < 0.05 were considered significant. Figures were prepared using Igor Pro 6 (WaveMetrics), Prism 7 (GraphPad, La Jolla, CA), Inkscape (GitLab), and Photoshop (Adobe).

### 2.7 Solutions for *in vitro* electrophysiology

We conducted slice preparation and recordings in ACSF solution contains (in mM): 120 NaCl, 2.5 KCl, 1.3 MgCl_2_, 10 glucose, 26 NaHCO_3_, 1.24 NaH_2_PO_4_, 4 CaCl_2_, 2 thiourea, 1 Na-L-ascorbate, 3 Na-pyruvate (pH 7.3-7.4 with 95% O_2_ and 5% CO_2_; 310-320 mOsm). We filled the pipette recordings with a Cs-methane-sulfonate-based solution contained (in mM): 125 Cs-methane-sulfonate, 11 KCl, 10 HEPES, 0.1 CaCl_2_, 1 EGTA, 5 Mg-ATP and 0.3 Na-GTP (pH adjusted to 7.2 with CsOH, 280 mOsm).

### 2.8 Perfusion and immunostaining

We deeply anesthetized mice with ketamine/xylazine (150/15 mg/kg, i.p.) and transcardially perfused them with PBS (Boston Bio, Product # BM300) followed by 10% formalin (Fisher, Product #SF100). We then removed the brains and stored them overnight in 10% formalin at 4°C. For cryoprotection, we submerged the brains in 25% sucrose (Sigma-Aldrich, Product # S0389) in phosphate-buffer saline (PBS) with sodium azide (Fisher, Product # 277l) for at least one day. Afterwards, we sectioned the brains at 30 μm (40 μm for CTb injections) in three series using a sliding microtome (Microm HM440E). The brain sections were collected into a cryoprotectant solution and stored at - 20°C for immunohistology.

To immunolabel CTb, we first rinsed sections three times with PBS (5 min each), and then incubated them in 0.3% hydrogen peroxide (Sigma-Aldrich, Product # H1009) for 30 min to quench endogenous peroxidase activity. After 3 additional washes in PBS, we incubated sections overnight in goat anti-CTb antiserum (List Biological, Product#: 703, Lot#: 7032A8, **Table 3**) diluted 1:20,000 in PBT with normal horse serum (NHS) and sodium azide at room temperature on a tissue shaker. The next day, we washed sections six times (5 min each) and then incubated them for 2 hours in biotinylated donkey anti-goat secondary antiserum (Jackson ImmunoResearch, Product#: 705-065-147, Lot#: 129472) diluted 1:500 in PBT with NHS. We then washed the sections three times with PBS (10 min each) and incubated them for 1 hour in avidin-biotin-complex (Vectastain ABC Elite Kit; Vector Laboratories). We washed sections four times in PBS (5 min each) and incubated them for 5 min in diaminobenzidine (DAB) solution to label CTb-containing neurons in brown.

**TABLE 3.**
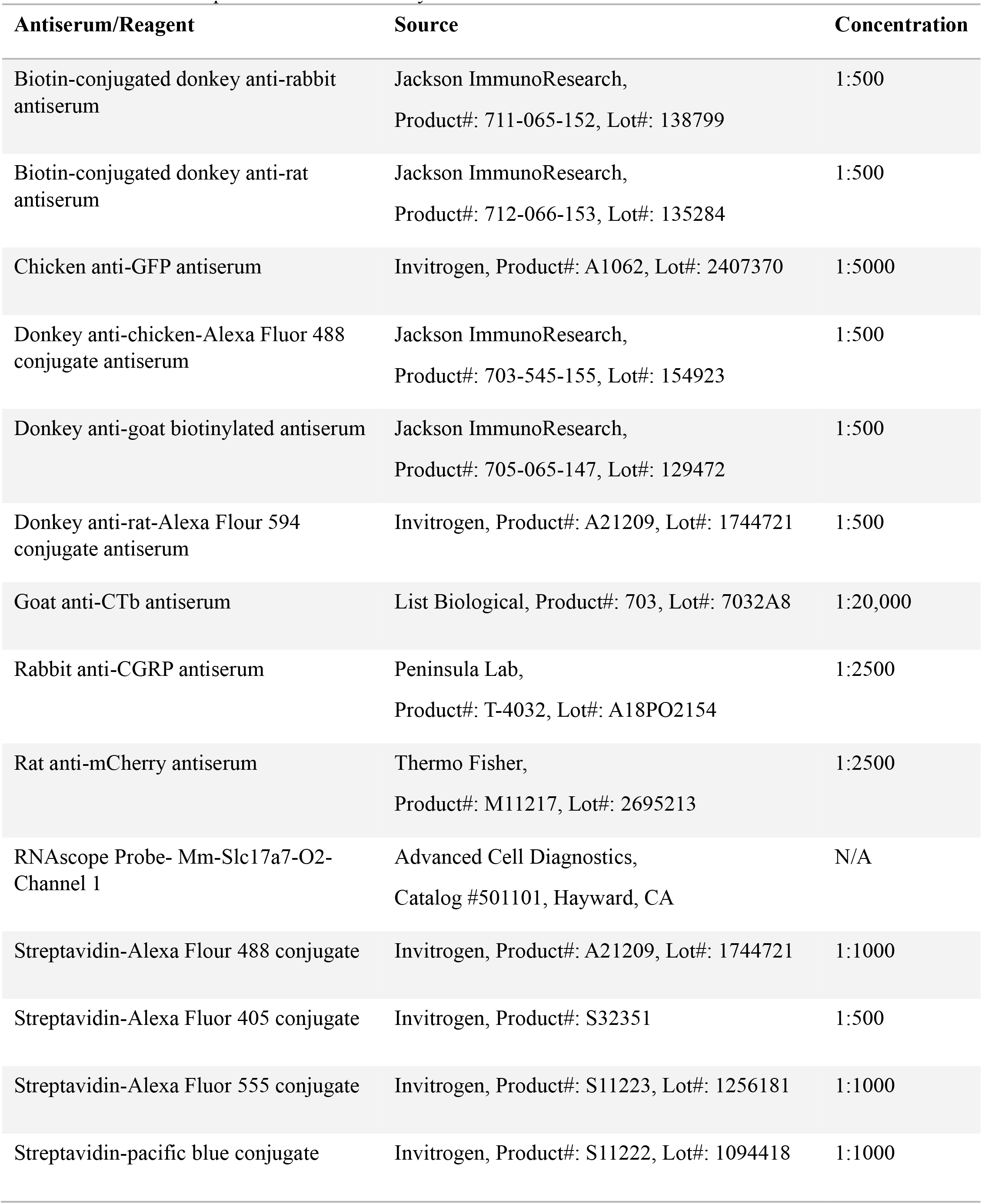
Antisera and probes used in this study.

**TABLE 4.**
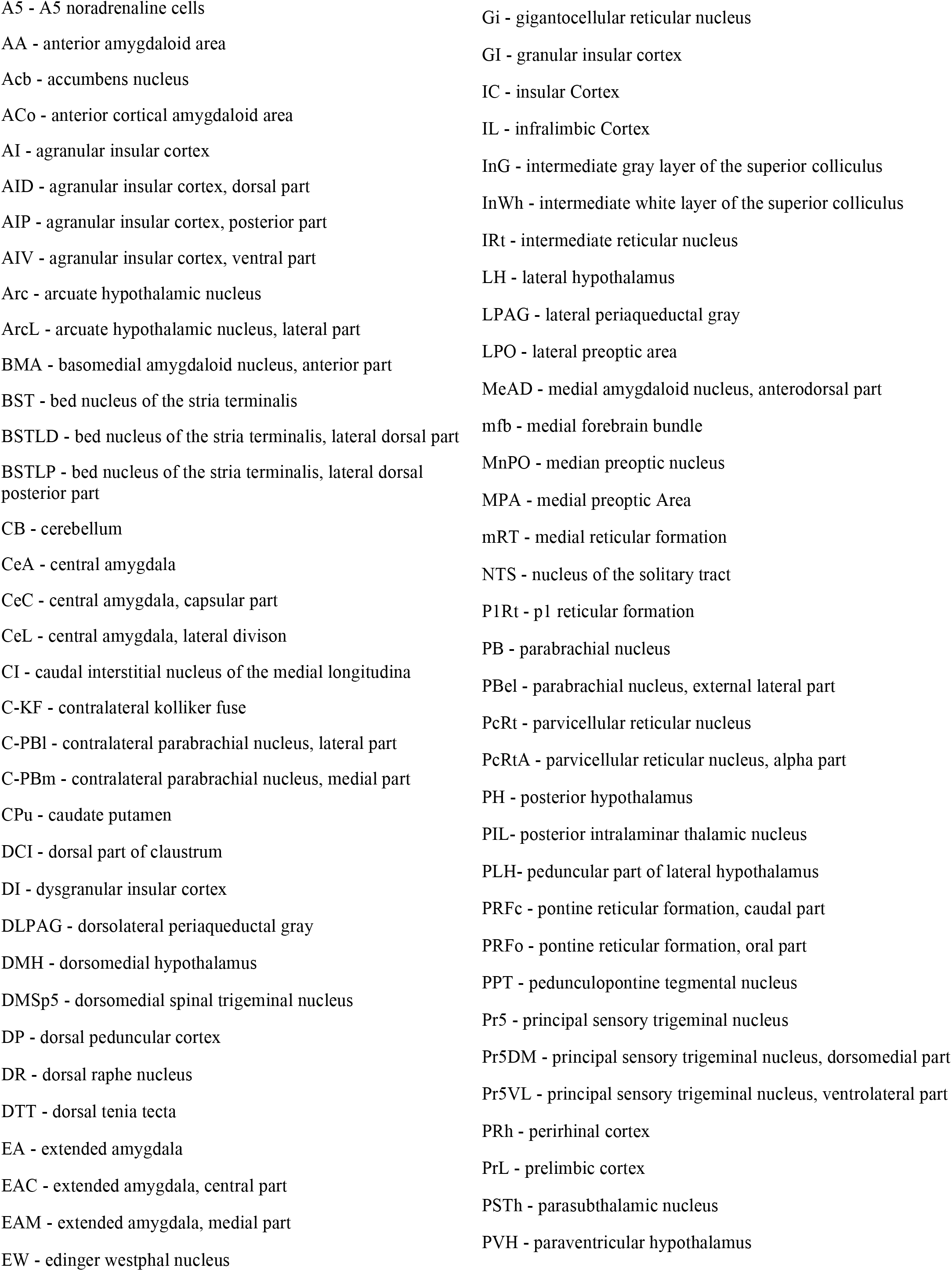
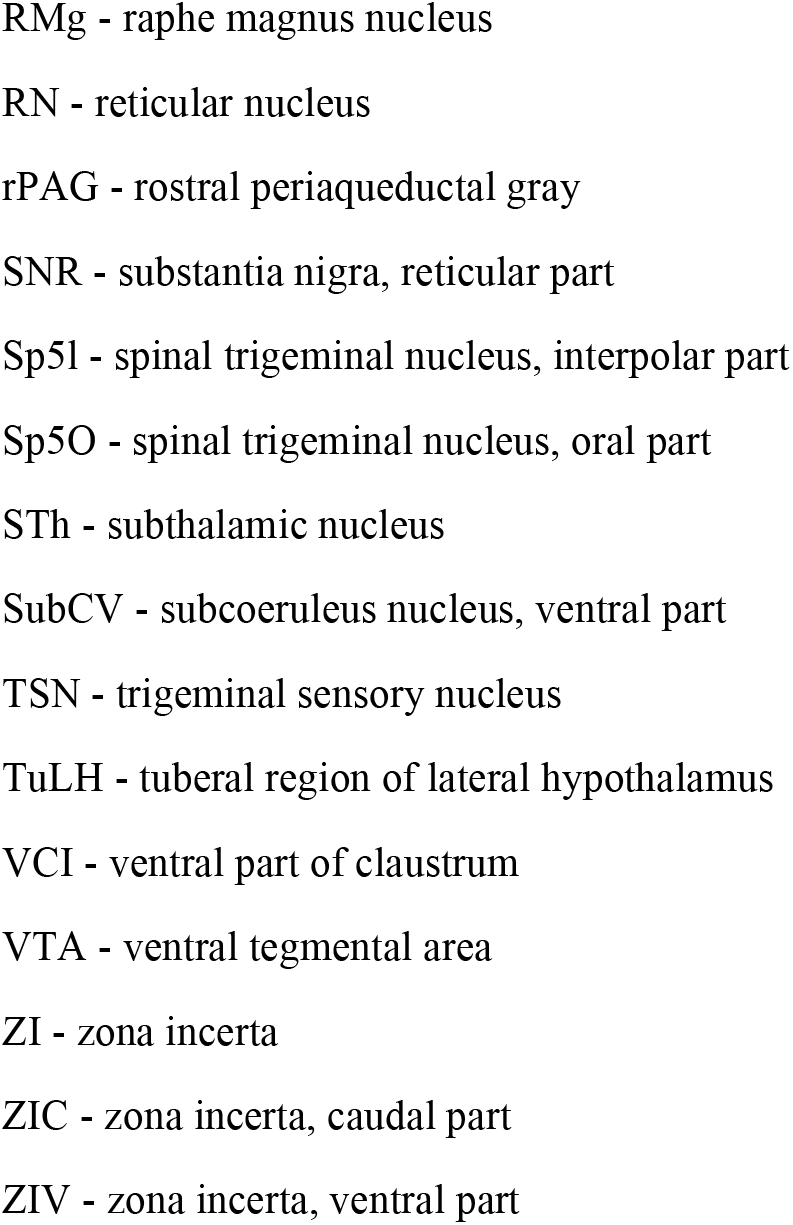
List of abbreviations.

For immunofluorescent labeling of CTb in *vGAT-* and *vGLUT2-reporter* mice, we washed sections three times with PBS (5 min each) and then incubated them overnight in goat anti-CTb antiserum (1:10000 in PBT with NHS and sodium azide) at room temperature. We then washed the sections six times with PBS (5 min each) and incubated them for 2 hours in biotinylated donkey anti-goat secondary antiserum (1:500) in PBT-NHS. After washing sections three times with PBS (10 min each), we incubated them for 1 hour in streptavidin-conjugated-Alexa Fluor 555 (Invitrogen, Product#: S11223, Lot#: 1256181), diluted 1:1000 in PBT with NHS.

For immunofluorescence labeling of CTb after *in situ* hybridization, sections were mounted on Superfrost Plus glass slides, washed three times with RNase-free PBS (5 minutes each) and then incubated overnight at 4°C in goat anti-CTb antiserum (diluted 1:5000 in RNase-free PBS). Subsequently, sections were washed three times with PBS (5 minutes each) and incubated for 2 hours in biotinylated donkey anti-goat secondary antiserum (diluted 1:500) in RNase-free PBS. After another three PBS washes, sections were incubated for 1 hour in streptavidin-conjugated Alexa Fluor 555, diluted 1:1000 in RNase-free PBS.

For the rabies experiments, we used fluorescence immunolabeling for TVA-mCherry and RV-GFP. We rinsed the brain sections three times with PBS (5 min each), and then incubated them overnight at room temperature in chicken anti-GFP antiserum (1:5000; Invitrogen, Product#: A1062, Lot#: 2407370) and rat anti-mCherry antiserum (1:2500; Thermo Fisher, Product#: M11217, Lot#: 2695213). The next day, we washed sections six times (5 min each) and incubated them for 2 hours in donkey anti-chicken-Alexa Flour 488 antiserum (Jackson ImmunoResearch, Product#: 703-545-155, Lot#: 154923) and biotinylated donkey anti-rat secondary antiserum (Jackson ImmunoResearch, Product#: 712-066-153, Lot#: 135284), diluted 1:500 in PBT with NHS. After washing sections three times (10 min each), we then incubated them for 1 hour in streptavidin-conjugated Alexa Fluor 555, diluted 1:1000 in PBT-NHS. For immunofluorescent labeling of hChR2-mCherry and CGRP, we washed the brain sections three times with PBS (5 min each), then incubated them overnight in rat anti-mCherry antiserum (1:2500; Thermo Fisher, Product#: M11217, Lot#: 2695213) and rabbit anti-CGRP antiserum (1:2500; Peninsula Lab, Product#: T-4032, Lot#: A18PO2154) at room temperature. The next day, we washed the sections six times (5 min each) and then incubated them for 2 hours in donkey anti-rat-Alexa Flour 594 antiserum (Invitrogen, Product#: A21209, Lot#: 1744721) and biotinylated donkey anti-rabbit secondary antiserum (Jackson ImmunoResearch, Product#: 711-065-152, Lot#: 138799) diluted 1:500 in PBT with NHS. We then washed the sections three times with PBS (10 min each) and incubated them for 1 hour in streptavidin-conjugated Alexa Flour 488 (Invitrogen, Product#: A21209, Lot#: 1744721) or streptavidin-pacific blue (Invitrogen, Product#: S11222, Lot#: 1094418, **Figure 2 (c)**), diluted 1:1000 in PBT with NHS.

After final washing of the brain sections three times in PBS, we mounted sections on glass slides. For DAB-stained sections, we air-dried the slides and dehydrated them through ascending concentrations of alcohols and then xylene before cover-slipping with Cytoseal (Fisher, Product# 8310-4). Sections with fluorescence labeling were air-dried and cover-slipped with Vectashield containing DAPI (Vector Lab Inc. Product # H-1500).

To map the injection sites for the CRACM experiments, we cut 30μm-thick slices containing the CeA after 3 nights fixation in 10% buffered formalin. To enhance the ChR2-EGFP signal, we immunolabeled the CeA neurons using chicken anti-GFP primary antiserum (Invitrogen, Product#: A1062, Lot#: 2407370), diluted 1:5000, and donkey anti-chicken-Alexa Fluor 488 secondary antiserum (Jackson, Product#: 703-545-155, Lot#: 154923), diluted 1:500. We wet-mounted the slices and scanned whole sections using a z-stack at 3-8 μm intervals. We viewed stacks of images using the OlyVIA slide scanner (Olympus) and software at final magnification of 10X to identify the ChR2 injection site. We used white matter tracts and anatomical landmarks to map the regions containing ChR2-EGFP neurons in the CeA onto template drawings using GIMP (version 10.02). We compiled the distribution of ChR2-EGFP expression in density plot using a Python script (www.python.org, version 2.7.5.0), representing the region of overlap of the maximum number of cases in red and fewest cases in blue. Immediately after the *in vitro* recordings, we fixed the recorded slices (250 μm-thick) in 10% buffered formalin, washed them, and incubated them overnight in streptavidin-conjugated Alexa Fluor 405 (1:500; Cat#: S32351; Invitrogen, Thermo Fisher Scientific Waltham, MA) to label the recorded PBel^*Calca*/CGRP^ neurons filled with biocytin (Anaclet et al., 2018; De Luca et al., 2022). We acquired images using a Leica Stellaris 5 confocal microscope at a final magnification of 63X.

### 2.9 *In Situ* Hybridization

We used RNAscope Multiplex Fluorescent Reagent Kit V2 (catalog #323100, Advanced Cell Diagnostics, Hayward, CA) for RNA scope *in situ* hybridization. Sections containing the IL and IC from wild-type mice were used for *vGLUT1 (Slc17a7)* mRNA labeling. For this, brains were fixed in formalin (10% buffered solution, overnight), cryoprotected (20% sucrose), cut into 20 μm sections and then mounted on Superfrost slides. All sections were washed with PBS and pretreated for 20 mins with hydrogen peroxide at room temperature. We then put sections in target retrieval reagent and steamed them at >99°C for 5 min. We dehydrated the sections in 90% alcohol and air-dried them for 5 min. We treated sections with protease reagent (Protease III) for 30 min at 40°C, and washed with sterile water. We then hybridized sections with probe for Slc17a7 (RNAscope Probe-Mm-Slc17a7-O2; catalog #501101, Advanced Cell Diagnostics, Hayward, CA) for 2 h at 40°C. We incubated sections in amplification reagents at 40°C (AMP1, AMP2 for 30 min each, and AMP3 for 15 min) followed by horseradish peroxidase-C1 (HRP-C1) at 40°C for 15 min. We incubated sections in TSA (Trichostatin A) plus Fluorescein (catalog #NEL741001, PerkinElmer, MA) at a 1:1000 concentration for 30 min to visualize vGLUT1 mRNA. In the final step, sections were subjected to HRP blocker for 15 min at 40°C. After each step, sections were washed with 1X wash buffer. We dried the slides and coverslipped them with Prolong Gold Antifade mountant (catalog # P36934, Invitrogen, Carlsbad, CA).

### 2.10 Microscopy and analysis

We used an Olympus VS120 slide-scanning microscope for imaging all slides. After reviewing the brain sections from whole-slide images in OlyVIA or VS-ASW software, we obtained high magnification images with a Leica Stellaris 5 confocal microscope. For dark-field imaging of brain sections, we used a Zeiss Axioplan 2 microscope equipped with a 20X objective. We used ImageJ software (version 1.52) to adjust the brightness, contrast, and apply filters to enhanced visibility of retrogradely labeled neurons in the brain sections. We then used the ‘Cell Counter’ plugin, accessible under Plugins > Analyze, for cell counting. After initializing the counter, we manually marked and counted each cell. Results and the annotated image were saved for future reference.

## 3 RESULTS

### 3.1 Inputs to the lateral PB region

We microinjected 3 nL of CTb into the lateral PB of 5 wild-type mice and immunolabeled brain sections for CTb 1 week later (**Figure 1**). We restricted our analysis to 4 mice with injection sites primarily limited to the lateral PB.

**FIGURE 1.**
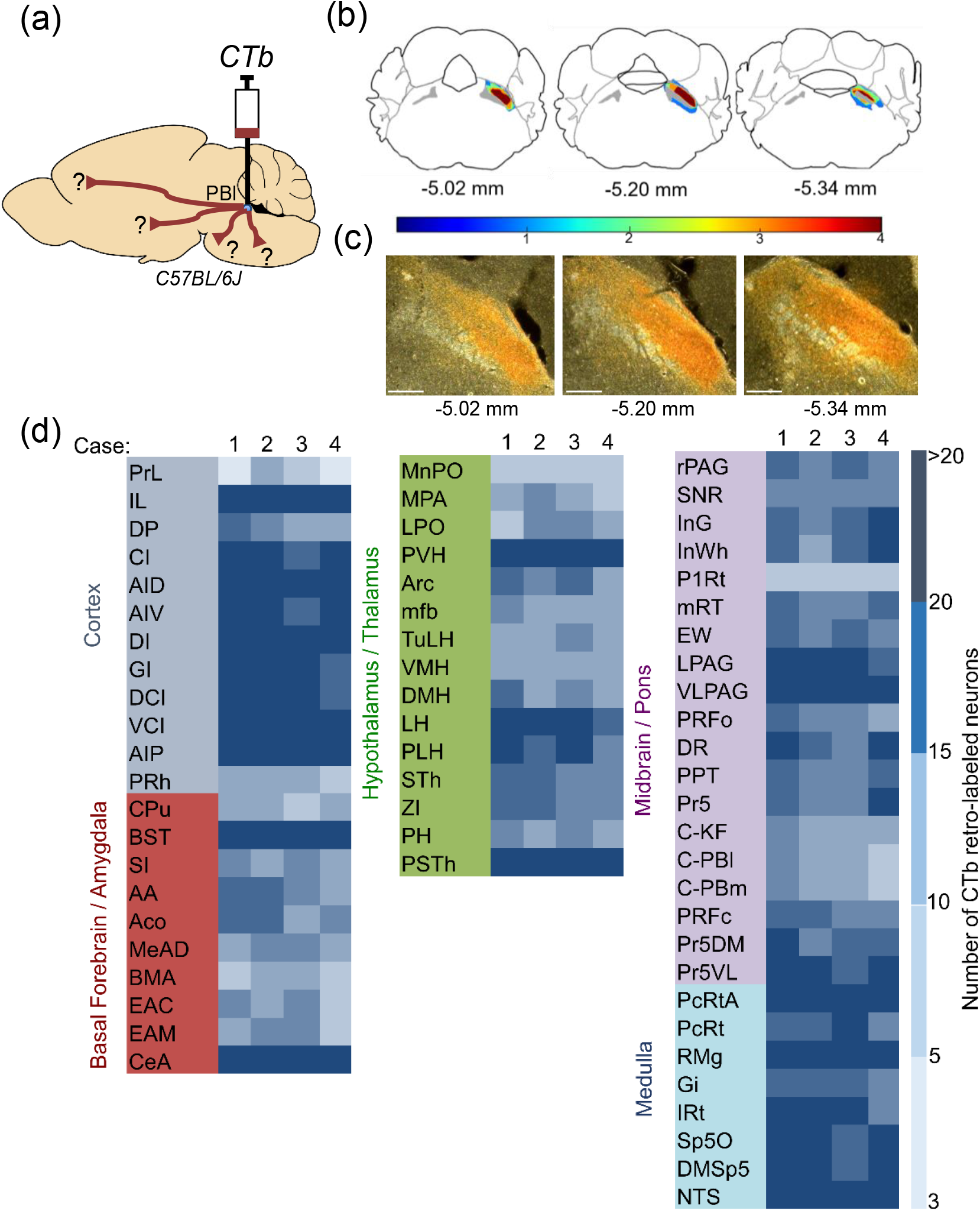
Retrograde labeling of inputs to the lateral PB using Cholera Toxin B subunit (CTb). (a) Schematic representation of CTb injections into the lateral PB of *C57BL/6J* mice. (b) Density plot of 4 injection sites in the lateral PB. (c) Representative CTb injection site in the lateral PB of wild-type mice. (d) Relative numbers of retrogradely labeled neurons in specific brain regions. Light blue: Regions with 3-5 CTb-labeled neurons; Dark blue: Regions with 20 or more CTb-labeled neurons. (d) Retrograde labeling with CTb was most abundant in the infralimbic and insular cortex, BST, CeA, PVH, LH, PSTh, LPAG, VLPAG, reticular nuclei, trigeminal nuclei, and NTS in all 4 mice. Scale bar in c is 100 μm. See Table 4 for list of abbreviations.

**FIGURE 2.**
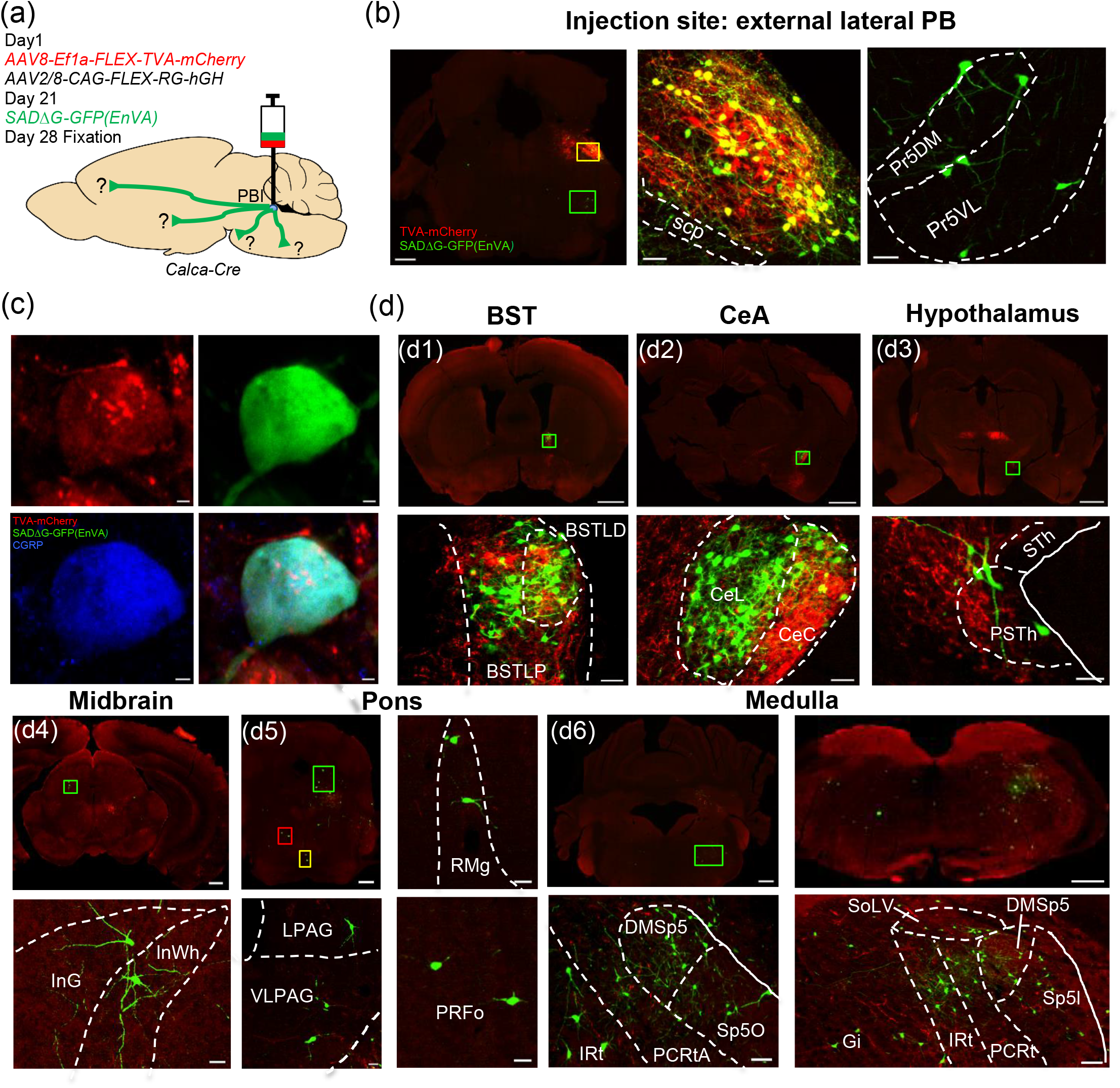
Retrograde labelling of monosynaptic inputs to the PB^*Calca*/CGRP^ neurons. (a) Schematic illustration of conditional rabies-based trans-synaptic retrograde tracing with injection of helper AAV followed 3 weeks later by conditional rabies. (b) Starter PB neurons (yellow) co-express TVA-mCherry (red) and rabies-GFP (green). (c) Most of these starter cells contain TVA-mCherry, GFP, and CGRP (d) The PB^*Calca*/CGRP^ neurons receive monosynaptic inputs from the BST, CeA, PSTh, and several nuclei in the brainstem. Scale bars: 1 mm in the top images of panels d1, d2, d3; 500 μm in the left image of panel b, and in the top images of panels d4 and d6, as well as the top-left image of panel d5; 100 μm in the middle and right images of panel b; 50 μm in the bottom images of panels d4 and d6, and the top-right image of panel d5; 5 μm in panels c.

CTb retrogradely labeled many neurons in ipsilateral forebrain and brainstem structures, including the IC, claustrum (Cl), BST, CeA, PVH, lateral hypothalamic area (LH), periaqueductal gray (PAG), and NTS. The same regions on the contralateral side had about 5-10% as many labeled cells.

#### 3.1.1 Cortex

We found numerous retrogradely labeled neurons in layer V of the IL, layer V of the agranular and dysgranular parts of the insular cortex (AI, DI), and the CI, medial to the IC, across its entire rostrocaudal axis. CTb labeling was moderate in neurons in the perirhinal cortex (PRh), dorsal peduncular cortex (DP), and slight in the prelimbic cortex (PrL).

#### 3.1.2 Basal Forebrain and Extended Amygdala

We observed numerous CTb-labeled neurons in the BST and amygdala. CTb labeled many neurons in the lateral division, dorsal and posterior parts of the BST, especially in the BSTLD. Both the lateral and medial divisions (CeL and CeM) of the CeA contained many labeled cells, and the rostral CeA contained a moderate number. In the extended amygdala between these regions, the SI contained scattered CTb-labeled cells.

#### 3.1.3 Hypothalamus

The PVH, LH, and PSTh showed extensive retrograde labeling, and moderate numbers of cell were labeled in the lateral preoptic area (LPO) medial preoptic areas (MPA), Arc, dorsomedial hypothalamic nucleus (DMH), subthalamic nucleus (STh), and zona incerta (ZI). In contrast, CTb-labeled neurons were less common in the median preoptic nucleus (MnPO) and ventromedial hypothalamic nucleus (VMH).

#### 3.1.4 Midbrain and Pons

The rostral part of the dorsolateral periaqueductal gray (rPAG) contained moderate numbers of CTb-labeled neurons, and the caudal part of the ventrolateral PAG contained many labeled cells. The caudal part of the pontine reticular formation (PRFc) and the principal sensory trigeminal nucleus exhibited numerous retrogradely labeled neurons. We also found moderate numbers of CTb-labeled neurons in the superior colliculus (intermediate gray (InG) and white (InWh) layers), dorsal raphe nucleus (DR), Edinger-Westphal nucleus (EW), mesencephalic reticular formation (mRT), and pedunculopontine tegmental nucleus (PPT).

#### 3.1.5 Medulla

The NTS contained many CTb-labeled neurons, with very dense labeling in the rostral part of the ventrolateral subnucleus. Labeled neurons were also abundant ventral to the NTS, in the spinal trigeminal nucleus (Sp5O, DMSp5), parvicellular reticular nuclei (PcRt, PcRtA), intermediate reticular nucleus (IRt), and gigantocellular reticular nucleus (Gi). While CTb-labeled neurons were present throughout the medullary reticular formation, their density was higher in the rostral parts. The raphe magnus nucleus (RMg) contained numerous retrogradely labeled neurons throughout its rostrocaudal extent.

### 3.2 Monosynaptic inputs to the PB^*Calca*/CGRP^ neurons

Many of the cells retrogradely labeled with CTb may innervate a variety of neurons in the lateral PB, so to identify neurons directly synapsing on the PB^*Calca*/CGRP^ neurons, we used conditional rabies virus tracing. First, we injected AAVs coding for Cre-dependent avian retroviral receptor (TVA) and rabies glycoprotein (RG) protein into the lateral PB of *Calca*-Cre mice to express TVA and RG selectively in *Calca/*CGRP-expressing neurons. Four weeks later, we injected genetically modified rabies virus SADΔG-GFP (EnvA) into the same region (**Figure 2**). A week later, we perfused the mice and immunostained their brains for mCherry to visualize TVA expression and GFP to identify rabies-infected neurons.

The helper virus injection sites were limited to the lateral PB, with slight extension into the medial PB. Most (72%) TVA-mCherry+ neurons showed CGRP-immunoreactivity in the external lateral PB (**Figure 2(c)**). In addition, about 60% of the TVA-mCherry-expressing neurons were also GFP-positive, indicating a relatively high efficiency of infection with SADΔG-GFP (EnvA). Nearly all these starter neurons (in yellow, **Figure 2**) were in the mid-rostrocaudal level of the external lateral PB, with a few scattered cells in the medial PB.

Primary afferents to the *Calca*/CGRP neurons of the external lateral PB were defined as GFP-positive but mCherry-negative (**Figures 3 and 4**). Though CTb labeled many inputs to the lateral PB from IL, IC and other cortical regions, GFP-positive, rabies-containing neurons were rare in the cortex. In contrast, the BSTLD and the CeL contained large numbers of first-order afferent neurons. Within the BSTLD and CeA, axons from the PB^*Calca*/CGRP^ neurons intermingled with many of these rabies-labeled neurons, suggesting reciprocal connections (**Figure 2(d)**). Surprisingly, the hypothalamus contained relatively few neurons, though the Arc, lateral hypothalamus, zona incerta, and PSTh contained moderate numbers.

**FIGURE 3.**
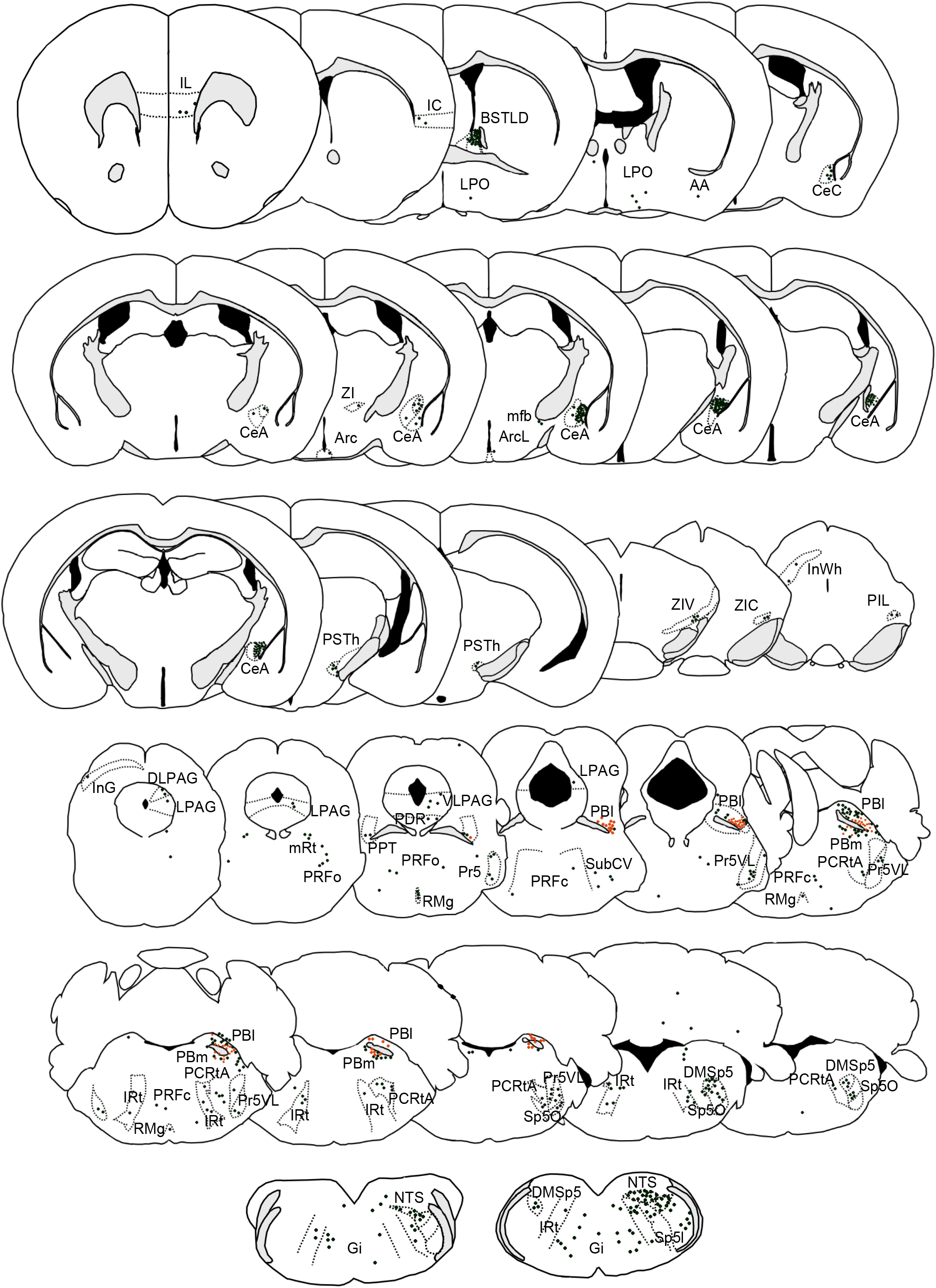
Coronal sections from a representative case illustrate the distribution of retrogradely labeled, monosynaptic inputs to the PB^*Calca*/CGRP^ neurons. Dark green dot: rabies-GFP + neurons; orange dot: starter neurons co-expressing TVA-mCherry and rabies-GFP.

**FIGURE 4.**
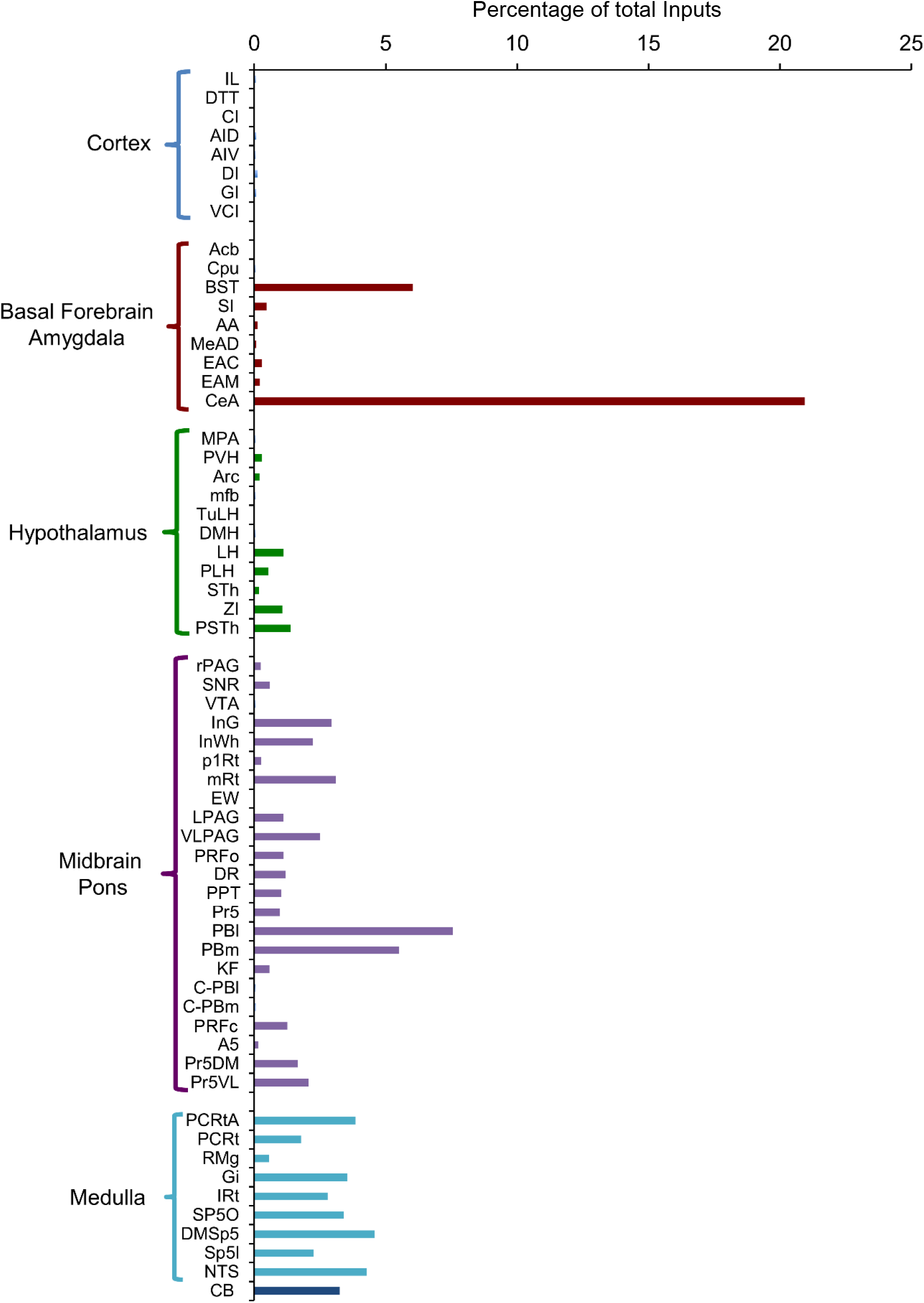
Relative proportions of rabies-labeled inputs to the PB^*Calca*/CGRP^ neurons. The extended amygdala (CeA, BST) contain over 25% of all labeled neurons, and the lateral PB contain local neurons innervating the PB ^*Calca*/CGRP^ neurons.

The upper brainstem showed moderate to large numbers of GFP-labeled cells in the superior colliculus (InG and InWh), the lateral and ventrolateral periaqueductal gray (LPAG, VLPAG), as well as in the pons in the RMg and the oral part of the pontine reticular formation (PRFo). Many single-labeled green neurons were present in the lateral and medial PB which are likely local afferents to the PB^*Calca*/CGRP^ neurons. In the medulla, GFP-expressing neurons were common in the reticular nucleus, primary and sensory trigeminal nucleus, Gi, and NTS (**Figure 2(d), 3, 4**).

### 3.3 Identifying GABAergic inputs to the lateral PB using retrograde tracing

To identify GABAergic inputs, we injected CTb into the lateral PB of *vGAT-GFP reporter* mice (**Figure 5**). One week later, we fixed the brains and immunostained the brain sections for CTb. The CTb injection sites were predominantly restricted to the lateral PB. GFP was present in the vast majority of CTb-labeled neurons within the medial, lateral, and capsular parts of the CeA (97%), the BSTLD (94%), and the SI (86%). Most CTb-labeled cells in the VLPAG (61%), and the Gi (61%) contained GFP. To a lesser extent, GFP expression was also observed in many neurons of the parvicellular reticular nucleus, alpha part (PCRtA, 20%) and the subcoeruleus nucleus, ventral part (SubCv,18%).

**FIGURE 5.**
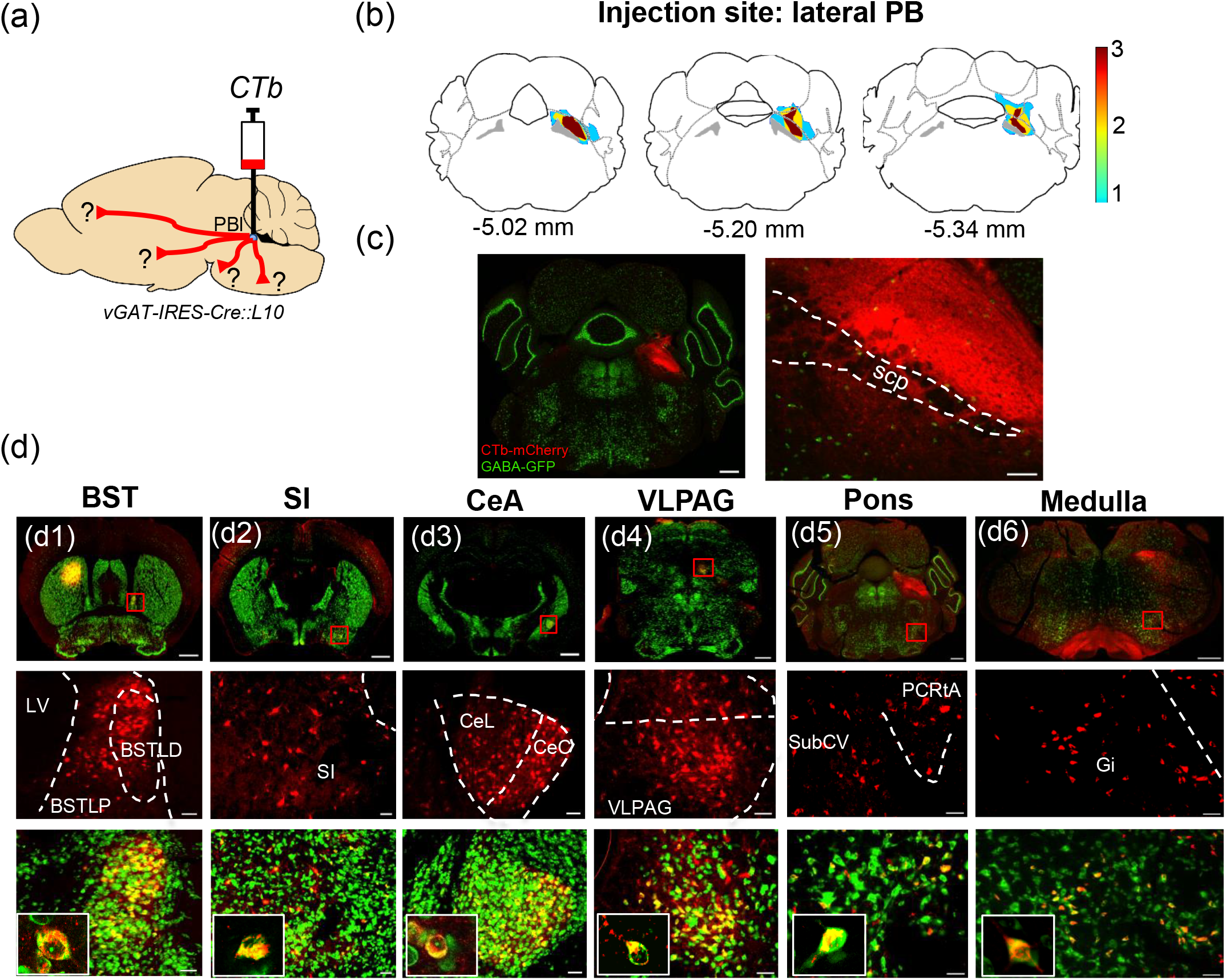
Retrograde labeling of GABAergic inputs to the lateral PB. (a) Schematic of CTb injection into the lateral PB of *vGAT-IRES-Cre::L10* reporter mice. (b) Density plot of injection sites in the PB. (c) Immunofluorescence labeling for CTb (red) and vGAT (green) in a typical injection site centered on the lateral PB. (d) Neurons co-expressing CTb and vGAT are common in the BST (94% of CTb-labeled cells contain vGAT), SI (86%), CeA (97%), vlPAG (61%), pons (38%), and medulla (61%). Scale bars: 1 mm in the top images of panels d1, d2, d3; 500 μm in the left image of panel c, and in the top images of panels d4, d5, d6; 100 μm in the right image of panel c, and in the middle and bottom images of panels d1, d2, d3, d4, d5, d6.

### 3.4 Identifying glutamatergic inputs to the lateral PB using retrograde tracing

To identify glutamatergic inputs to the lateral PB nucleus, we injected CTb into the lateral PB of *vGLUT2-GFP reporter* mice (**Figure 6**). One week later, we fixed the brains and immunostained brain sections for CTb. The CTb injection sites were predominantly restricted to the lateral PB.

**FIGURE 6.**
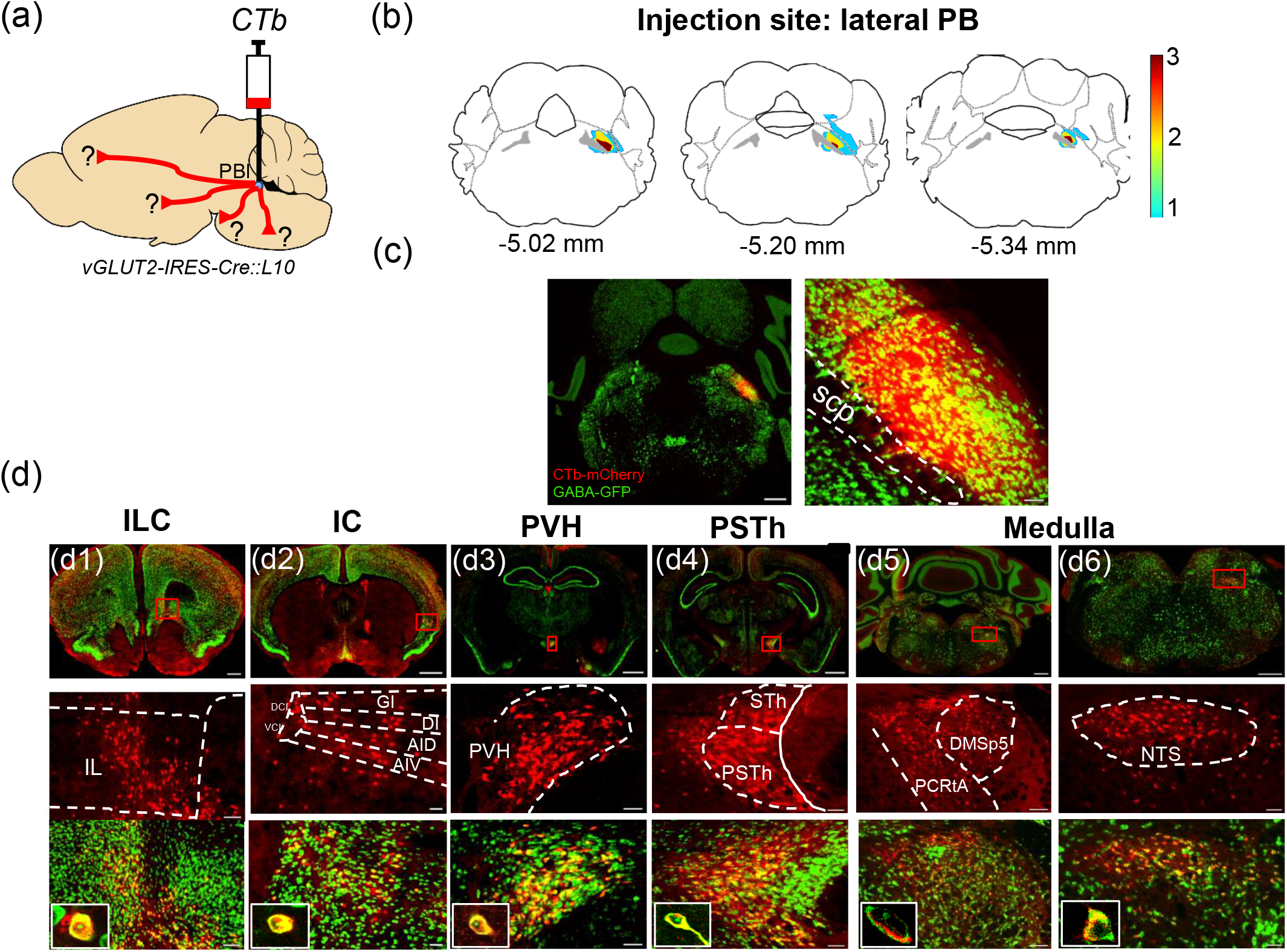
Retrograde labeling of glutamatergic inputs to the lateral PB. (a) Schematic of CTb injection into lateral PB of *vGLUT2-IRES-Cre::L10* reporter mice. (b) Density plot of injection sites in the PB. (c) Immunofluorescence labeling for CTb (red) and vGAT (green) in a typical injection site centered on the lateral PB. (d) CTb was common in glutamatergic neurons of the ILC (85% of CTb-labeled cells contained GFP), IC (63%), PVH (94%), PSTh (90%), and medulla (80%). Scale bars: 1 mm in the top images of panels d2, d3, d4; 500 μm in the left image of panel c, and in the top images of panels d1, d5, d6; 100 μm in the right image of panel c, and in the middle and bottom images of panels d1, d2, d3, d4, d5, d6.

GFP was present in most CTb-labeled neurons in the IL (85%), IC (63%), PVH (94%), PSTh (90%), and in the medulla, the PCRtA (75%), dorsomedial spinal trigeminal nucleus (DMSp5, 75%), and NTS (83%). Cortical neurons primarily express vGLUT2 during development and express vGLUT1 in adults (Boulland et al., 2004a; Nakamura et al., 2005; Berry et al., 2012). In situ hybridization for *vGLUT1 (Slc17a7)* mRNA revealed that the CTb-labeled neurons in the IL and IC express vGLUT1 mRNA (**Figure 7**).

**FIGURE 7.**
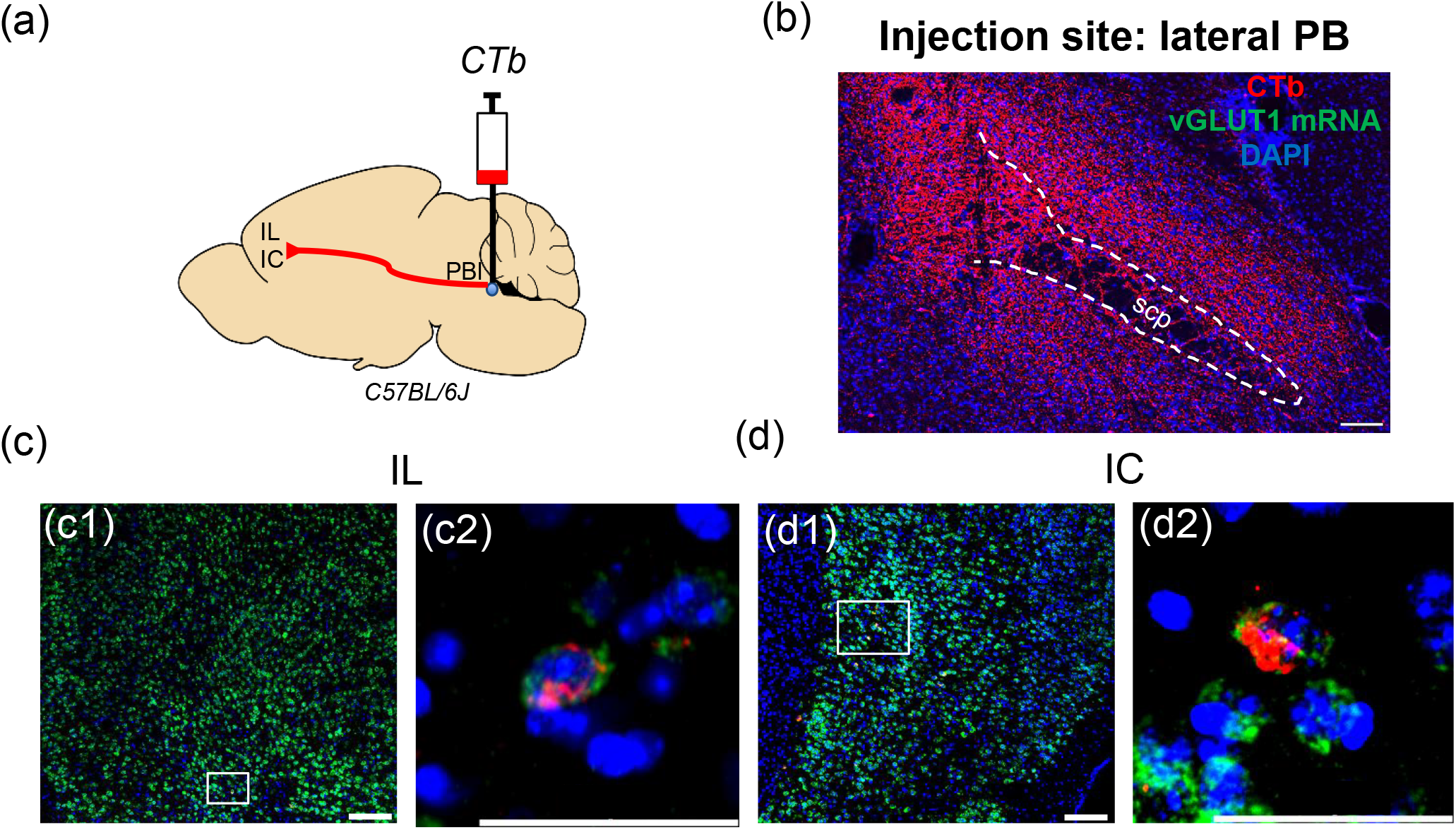
Retrogradely labeled neurons from lateral PB in infralimbic (IL) and insular cortex (IC) express vGLUT1 (*Slc17a7*) mRNA. (a) Schematic of CTb injection into lateral PB of wild-type mice. (b) CTb injection site in the PB. (c, d) vGLUT1 (*Slc17a7*) mRNA (green) co-localizes with CTb immunoreactivity (red) in IL and IC. Scale bars:100 μm in the image of panels b, c1, d1; 50 μm in the image of panels c2, d2.

### 3.5 Anterograde tracing of GABAergic axon terminals onto PB*Calca*/CGRP neurons

To determine if GABAergic neurons closely appose PB^*Calca*/CGRP^ neurons, we injected *AAV8-Ef1a-DIO-hChR2-mCherry* into several brain regions of *vGAT-Cre* mice (**Figure 8**), we then immunolabeled sections for mCherry and CGRP, and examined 1 μm optical sections of the PB using confocal microscopy.

**FIGURE 8.**
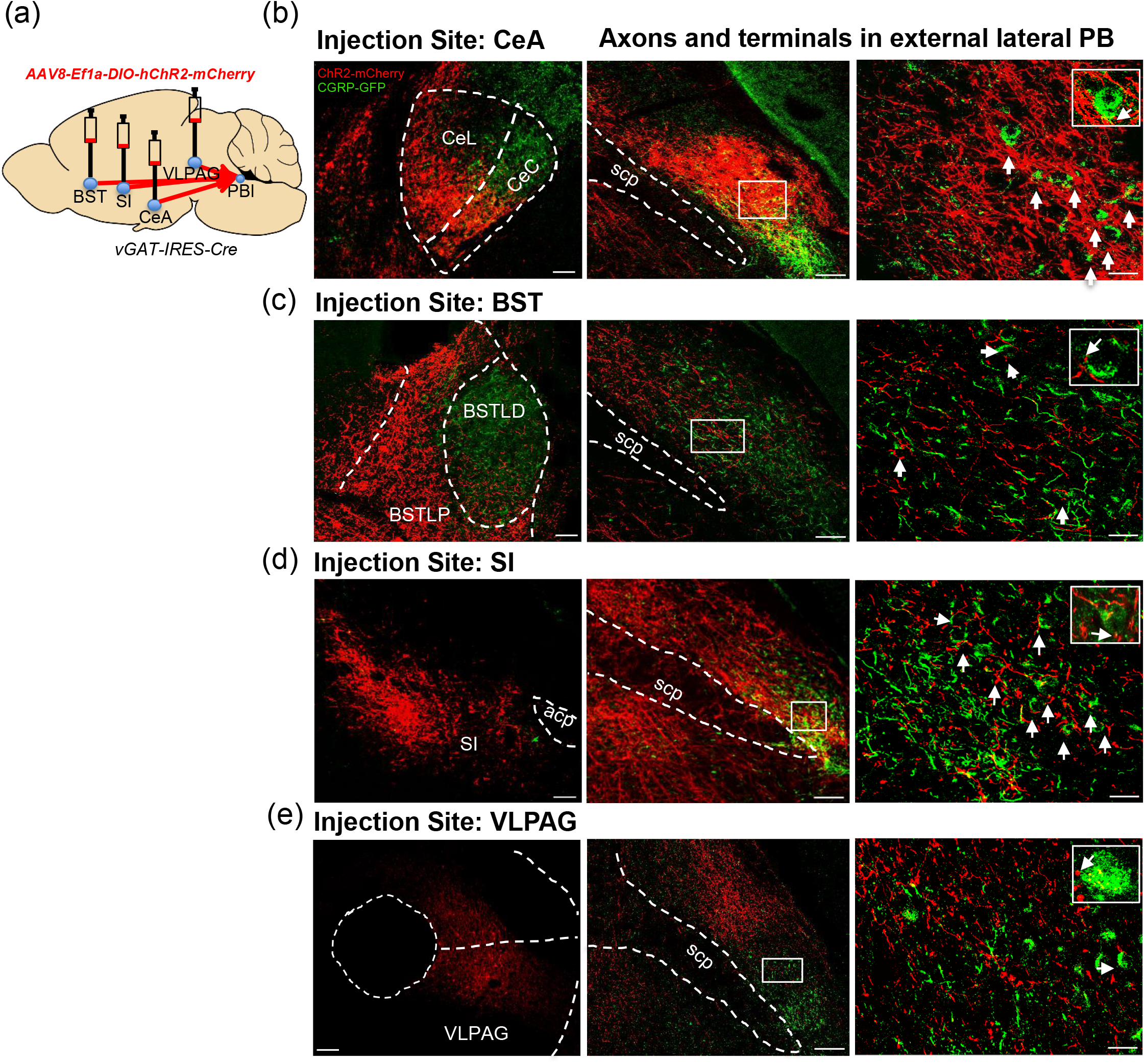
GABAergic nerve terminals and axons closely appose PBel^*Calca*/CGRP^ neurons. (a) *AAV8-DIO-hChR2-mCherry* was microinjected into the CeA, BST, SI, and VLPAG of vGAT-IRES-Cre mice. (b-e) Immunolabeling for ChR2-mCherry (red) and *Calca*/CGRP (green) at injection sites and in the PBel. GABAergic axon terminals from the CeA, BST, SI, vlPAG closely appose the PBel^*Calca*/CGRP^ neurons (white arrowheads). Scale bars: 100 μm in left and center images of panels b, c, d, e; 50 μm in right images of panels b, c, d, e.

In *vGAT-Cre* mice, the injection of *AAV8-EF1a-DIO-hChR2-mCherry* into the CeA densely labeled axon terminals in the rostral part of the lateral PB, and these terminals closely apposed the soma and dendrites of many PB^*Calca*/CGRP^ neurons. Injections into the BST resulted in modest labeling of axon terminals in the rostral part of the lateral PB, accompanied by a moderate number of close appositions on PB^*Calca*/CGRP^ neurons. However, these injections surrounded but only partially entered the BSTLD, and they may underrepresent the strength of this pathway. Injection into the SI produced moderate labeling in the same region but with more close appositions on the PB^*Calca*/CGRP^ neurons. Finally, injections in the VLPAG labeled a few close appositions on the soma and dendrites of the PB^*Calca*/CGRP^ neurons in the dorsal and lateral parts of PB. The ChR2-mCherry projections mainly labeled ipsilateral axonal terminals with sparse axons contralaterally.

### 3.6 Anterograde tracing of glutamatergic axon terminals onto PB^*Calca*/CGRP^ neurons

To identify glutamatergic inputs, we injected *AAV8-Ef1a-DIO-ChR2-mCherry* into several brain regions of *vGLUT2-Cre* mice (**Figures 9**). AAV injections in the PVH, PSTh, and NTS densely labeled axon terminals in the lateral PB, with numerous close appositions on the soma and dendrites of PB^*Calca*/CGRP^ neurons. As with all inputs, these projections were overwhelmingly ipsilateral.

**FIGURE 9.**
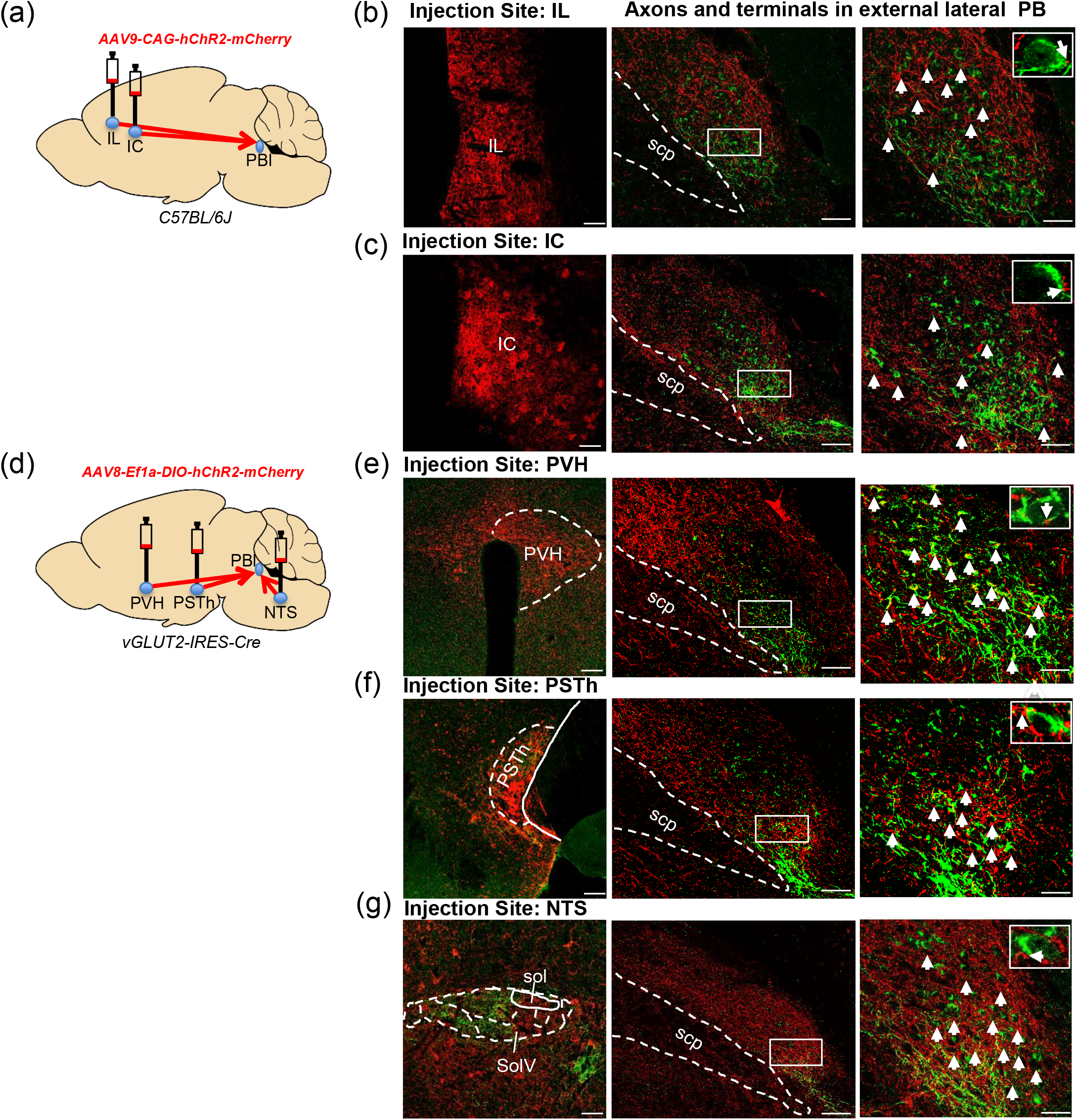
Glutamatergic nerve terminals and axons closely appose PBel^*Calca*/CGRP^ neurons. (a) *AAV9-hChR2-mCherry* was microinjected into the IL and IC of *C57BL/6J* mice. (b-c) Immunolabeling for ChR2-mCherry (red) and *Calca*/CGRP (green) at injection sites and in the PBel. (d) *AAV8-DIO-hChR2-mCherry* was microinjected into the PVH, PSTh, and NTS of *vGLUT2-IRES-Cre* mice. (e-g) Immunolabeling for ChR2-mCherry and CGRP at injection sites and in the PBeL. Glutamatergic axon terminals from the IL, IC, PVH, PSTh and NTS closely appose PBel^*Calca*/CGRP^ neurons (white arrowheads). Scale bars: 100 μm in left and center images of panels b, c, e, f, g; 50 μm in right images of panels b, c, e, f, g.

As the vGLUT2 promoter is generally inactive in the adult cortex, we labeled projections from the insular and infralimbic cortex of wild-type mice using non-conditional *AAV9-CAG-hChR2-mCherry*. This method labeled a moderate number of axon terminals in the lateral PB, with numerous close appositions on the soma and dendrites of CGRP neurons.

### 3.7 CeA^GABA^ neurons inhibit PBel^*Calca*/CGRP^ neurons

The retrograde and anterograde tracing experiments suggest that the CeA^GABA^ neurons innervate the PB^*Calca*/CGRP^ neurons, and to determine if these cells directly inhibit the PBel^*Calca/*CGRP^ neurons, we used ChR2-assisted circuit mapping (CRACM). Using *vGAT-flp::CGRP/Calca-Cre* mice, we injected the CeA with *AAVd/j-Ef1a-fDIO-hChR2-EGFP* and the PBel with *AAV8-DIO-mCherry*, thus expressing ChR2 in the CeA^GABA^ neurons and mCherry in the PBel^*Calca/CGRP*^ neurons, respectively. Using *ex vivo* slices of the PB, we then made whole cell recordings of mCherry-labeled PBel^*Calca*/CGRP^ neurons while photostimulating the CeA^GABA^ axons and terminals (0.1Hz, 10ms light pulses) in the PB. Light pulses produced opto-evoked inhibitory postsynaptic currents (oIPSCs) in 42% of the recorded PBel^*Calca*/CGRP^ neurons (11/26 neurons) (**Figure 10**). Latency and amplitude of the oIPSCs were 8.6 ± 0.94 ms and 18.31 ± 3.61 pA, respectively. Bicuculline (20 μM, *n* = 5 neurons) abolished the oIPSCs, indicating that this inhibition is mediated by GABA_A_ receptors.

**FIGURE 10.**
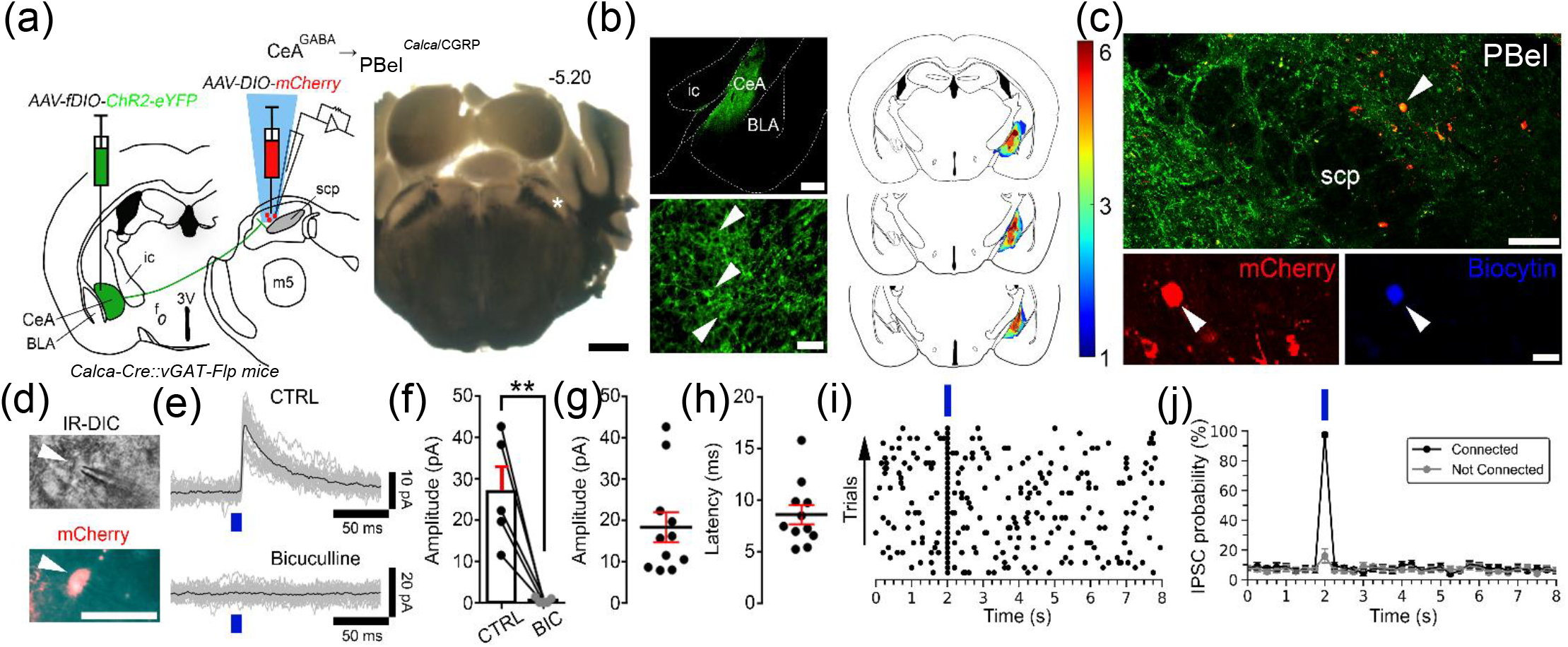
GABAergic CeA neurons (CeA^GABA^) directly inhibit PBel^*Calca*/CGRP^ neurons. (a) Experimental design for testing CeA^GABA^ → PBel^*Calca*/CGRP^ connectivity in brain slices. We injected *AAV-fDIO-ChR2-eYFP* into the CeA and *AAV-DIO-mCherry* into the lateral/external lateral PB of *Calca-Cre::vGAT-Flp* mice. We recorded from PBel^*Calca*/CGRP^ neurons labeled with mCherry while photostimulating CeA^GABA^ axon terminals in the PB. On the *right*, is an example of a recorded slice (−5.20 mm from bregma); (*) marks the location where *in vitro* recordings were conducted; scale bar: 1mm. (b) Photomicrographs of ChR2-eYFP expression in the CeA at low (*top*) and higher magnification (*bottom*, white arrows pointing at ChR2-eYFP expressing neurons within the CeA; scale bars: 500, 50 μm). On the *right*, density plots of the CeA injection sites (*n*=6). (c) An example of a recorded PBel^*Calca*/CGRP^ neuron labeled with mCherry (red) (*top*, low magnification) and *post hoc* labeled for biocytin with streptavidin-conjugated Alexa Fluor 405 (blue) at high magnification (*bottom*; white arrow). Scale bars: 100 and 20 μm. (d) IR-DIC and fluorescent visualization of PBel^*Calca*/CGRP^ neurons in brain slices. Scale bar: 50 μm. (e) Photostimulation of CeA^GABA^ terminals expressing ChR2-eYFP produce opto-evoked inhibitory postsynaptic currents (oIPSCs) in the PBel^*Calca*/CGRP^ neurons that are mediated by the GABA_A_ receptors as they are blocked by bicuculline (BIC 20 μM; GABA_A_ receptor antagonist). 30 individual oIPSCs (grey) and average IPSC (black) (blue bar represents a 10-ms light pulse). (f) oIPSC amplitude under control conditions (CTRL) and with BIC in 5 recorded PBel^*Calca*/CGRP^ neurons (*paired t-test*, one-tailed; **, *p*=0.0053). (g-h) Mean oIPSC amplitude and latency in PBel^*Calca*/CGRP^ neurons with photostimulation of the CeA^GABA^ input (*n*=11; means ± SEM). (i) Raster plot of GABAergic IPSCs in a representative PBel^CGRP^ neuron, during and after photostimulation of the CeA^GABA^ input (bin duration: 50ms). (j) Average IPSC probability in PBel^*Calca*/CGRP^ neurons that responded to the photostimulation of the CeA^GABA^ input (black; *n*=11) and those that did not respond (grey; *n*=15).

## 4 DISCUSSION

The PB^*Calca*/CGRP^ neurons are essential for responding to pain, hypercarbia, and a variety of other interoceptive signals (Carter et al., 2013, 2015; Han et al., 2015; Saper, 2016; Campos et al., 2017; Kaur et al., 2017; Palmiter, 2018), yet relatively little is known about their neuronal inputs. Using a variety of techniques, we identified afferents to the PB^*Calca*/CGRP^ neurons in mice (summarized in **Figure 11**). Retrograde tracing with CTb showed that the lateral PB receives inputs from the IL, IC, BST, CeA, PVH, LH, and NTS. Using conditional retrograde tracing with rabies virus, we found that the PB^*Calca*/CGRP^ neurons are heavily innervated by the BSTLD and CeL and moderately highlighting the internal connectivity within the PB. Using *vGAT-* and *vGluT2-reporter* mice and *in situ* hybridization, we found that inputs from the BST, CeA, SI, and the VLPAG are GABAergic whereas those from the IL, IC, PVH, PSTh, NTS and several other regions in the medulla are glutamatergic. Last as a test of functionality, we established that the GABAergic CeA neurons directly inhibit the PBel^*Calca*/CGRP^ neurons via GABA_A_ receptors. Collectively, these experiments shed light on the many neuronal systems that can influence the activity of the PB^*Calca*/CGRP^ neurons.

**FIGURE 11.**
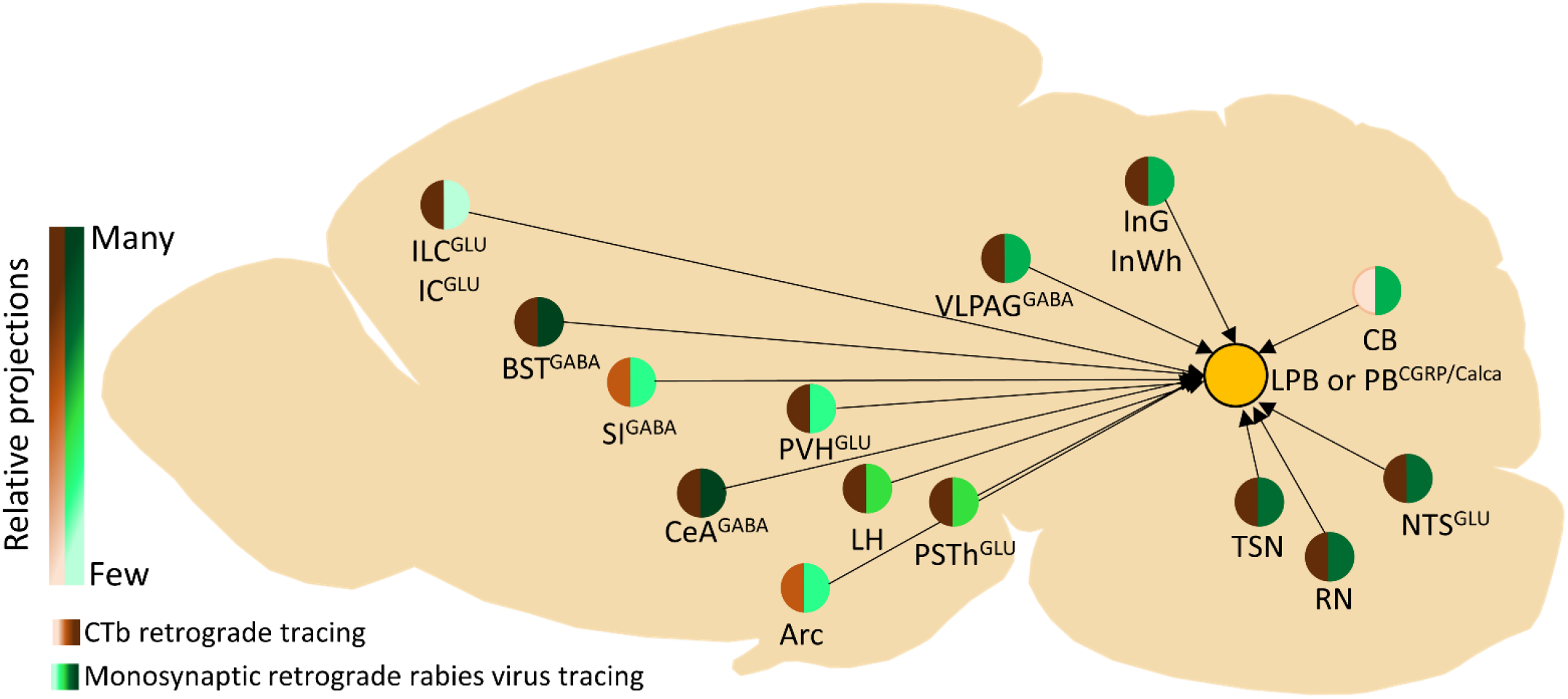
Summary of major inputs to the PB^*Calca*/CGRP^ neurons. In general, many regions retrogradely labeled with CTb also show *Calca*/CGRP-specific labeling with rabies. The PB^*Calca*/CGRP^ neurons receive strong inputs from GABAergic BST and CeA neurons and moderate GABAergic inputs from the SI and vlPAG. They also receive strong or moderate inputs from glutamatergic neurons of the PVH, PSTh, and NTS. Some regions such as the IL, IC SI, Arc and PVH show strong or moderate labeling with CTb but little labeling with rabies, suggesting these regions may innervate non-*Calca*/CGRP PB neurons.

### 4.1 Comparison with prior studies

Prior research using conventional retrograde and anterograde tracers in rats showed that the lateral PB receives inputs from the IL, IC, BSTLD, CeA, PAG, NTS, spinal dorsal horn and several other regions (Herbert et al., 1990; Moga et al., 1990; Hurley et al., 1991; Grady et al., 2020). Tokita and colleagues showed a similar pattern in mice, with perhaps fewer retrogradely labeled neurons in the MPA of mice than in rats (Tokita et al., 2009), and our CTb experiments in mice show a similar pattern though we did not find retrogradely labeled neurons in the ventral tegmental area. Overall, the afferents to the lateral PB appear similar in mice and rats.

These traditional tracing methods are very effective in identifying inputs to a PB region, but the PB contains a variety of distinct neuronal populations, and the inputs to the *Calca*/CGRP neurons are not well understood. Like most PB neurons, the *Calca*/CGRP neurons are glutamatergic and express vGLUT2, but they are distinct from other cell types. *Calca*/CGRP neurons are most easily identified in the PBel with high *Calca*/CGRP expression, yet *Calca*/CGRP is also expressed in medial PB and KF neurons with much less CGRP immunoreactivity (Kaur et al., 2013, 2017; Chen innervated by the PAG, superior colliculus, and additional brainstem regions. In addition, rabies labeled many neurons within the lateral and medial PB which are likely local afferents to the PB^*Calca*/CGRP^ cells, et al., 2018; Huang et al., 2021b; Kaur et al., 2023). In contrast, neurons containing dynorphin are abundant in the central, lateral, and dorsal lateral PB, but they do not overlap with the *Calca*/CGRP expressing neurons in the PBel (Miller et al., 2012; Geerling et al., 2016; Lee et al., 2019; Huang et al., 2021a). The transcription factor FoxP2 is expressed by many medial and dorsal PB neurons, including some of the dynorphin cells, but these are also anatomically distinct from the PB^*Calca*/CGRP^ cells (Miller et al., 2012; Geerling et al., 2016; Verstegen et al., 2017; Huang et al., 2021a; b; Kaur and Saper, 2021; Kaur et al., 2023). Additional markers such as *Tacr1* and *Lmx1b* also help distinguish these populations. (Miller et al., 2012; Huang et al., 2021b) (Barik et al., 2021).

Recent RNAseq studies have significantly advanced our understanding of PB neuron populations. For instance, Nardone and colleagues described 10 distinct subtypes of *Calca*/CGRP neurons, each characterized by specific marker genes (Nardone et al., 2024). These subtypes cluster by gene expression and collectively represent one of the largest neuronal groups in the PB, identified as *Calca*+/*Il20ra*+. Among these clusters, clusters #4 (marked by *Slc6a2+*) and #8 (marked by *Qrfpr+*) also show a distinct spatial distribution: cluster #4 resides in the apical/dorsal part of the main *Calca*+/*Il20ra*+ group, while cluster #8 is located more ventrally. The unique transcriptional profiles and spatial localizations of these *Calca*/CGRP neuron subsets raise intriguing questions about their potentially distinct functions and connectivity patterns (Pauli et al., 2022; Nardone et al., 2024).

The extended amygdala, especially parts of the BST and CeA, is the largest input to the PB^*Calca*/CGRP^ neurons. Conventional retrograde and anterograde tracing identified very strong inputs from the BSTLD and CeA in a pattern similar to prior reports (Moga et al., 1990; Tokita et al., 2009). Our findings build upon this by showing that these regions send terminals which closely appose the PB^*Calca*/CGRP^ neurons, and retrograde rabies virus tracing shows that these neurons innervate PB^*Calca*/CGRP^ neurons. Importantly, the CRACM experiments demonstrate that GABAergic neurons of the CeA directly inhibit PB^*Calca*/CGRP^ neurons via GABA_A_ signaling. The BSTLD and CeL have strikingly similar histology, neurochemistry and connections (McDonald, 1982, 1983; Roberts et al., 1982; Schwaber et al., 1982; Woodhams et al., 1983; Paxinos, 2004; Shin et al., 2008), and as they together account for more than 25% of all rabies-labeled cells, more work is needed to establish just how they impact the functions of the PB^*Calca*/CGRP^ neurons. CTb injections into the lateral PB also labeled a moderate number of neurons in the SI. These might be scattered, extended amygdala neurons akin to those in the BST and CeA (Paxinos, 2004). In addition, these may innervate non-Calca cells of the lateral PB as rabies labeled few cells in the SI, and anterograde tracing of GABAergic SI neurons showed only moderate axonal labeling near PBel^*Calca*/CGRP^ neurons.

In line with prior studies (Saper, 1982; Moga et al., 1990; Hurley et al., 1991; Tokita et al., 2009), we found that CTb injections into the lateral PB retrogradely labeled neurons primarily in layer V of the IL and AI, and anterograde tracing from these regions labeled many close appositions on the PBel ^*Calca*/CGRP^ neurons. However, monosynaptic tracing with rabies in *Calca-Cre* mice revealed few infected cells in the cortex, so despite the appearance of close appositions, these cortical regions may innervate other PBel cell types. Alternatively, they may directly innervate the PBel^*Calca*/CGRP^ neurons, but conditional rabies fails to label these cells (see Limitations section below); this idea could be further tested with functional methods such as CRACM.

Additional regions thought to be major inputs to the PB^*Calca*/CGRP^ neurons may actually innervate other cells in the lateral PB. For example, CTb labeled many glutamatergic neurons in the PVH and PSTh similar to prior reports (Moga et al., 1990; Tokita et al., 2009), but rabies tracing into PB^*Calca*/CGRP^ neurons labeled few neurons in these areas (though see Limitations section below). Similarly, CTb labeled many GABAergic neurons of the VLPAG region, but rabies showed few cells in this area, and anterograde tracing of GABAergic VLPAG neurons shows the terminal labeling is mainly dorsal to the PB^*Calca*/CGRP^ neuron field.

The PB receives many projections from the medulla, including the reticular formation, spinal trigeminal nucleus, and NTS (Norgren and Leonard, 1971; Herbert et al., 1990; Feil and Herbert, 1995; Tokita et al., 2009). CTb injections into the lateral PB strongly label these regions, and conditional rabies virus tracing labeled many neurons in these hindbrain regions, suggesting direct projections to the PB^*Calca*/CGRP^ neurons. The lateral PB also receives projections from the retrotrapezoid nucleus (RTN) in rats (Bochorishvili et al., 2012), but we did not observe retrogradely labeled neurons in the RTN of mice. Pain signals from the face are relayed through both a monosynaptic route from the trigeminal ganglia (TG) directly to the lateral PB, including PB^*Calca*/CGRP^ cells, and an indirect disynaptic route from TG to spinal trigeminal nucleus caudalis (Sp5C) to lateral PB (Rodriguez et al., 2017). In contrast, nociceptive signals from the rest of the body are relayed from dorsal root ganglia through the spinal dorsal horn to the lateral PB (Rodriguez et al., 2017). Inputs from the NTS are especially strong though we found more inputs from the rostral ventral NTS rather than the caudal NTS as described previously (Herbert et al., 1990; Tokita et al., 2009).

Many regions that innervate the PB^*Calca*/CGRP^ neurons receive reciprocal projections from the PB^*Calca*/CGRP^ neurons. Using conditional anterograde tracing in *Calca-Cre* mice, Huang and colleagues showed that the PB^*Calca*/CGRP^ neurons moderately or strongly innervate the CeA, BST, basal forebrain, PSTh, PAG, and spinal trigeminal nucleus (Huang et al., 2021b), and in our rabies experiments, all these regions contained inputs to the PB^*Calca*/CGRP^ neurons. This reciprocal pattern is quite apparent in our Figure 2d in which TVA-mCherry containing axons from PB^*Calca*/CGRP^ neurons intermingle with or are close to rabies-containing CeA and BST neurons that innervate the PB^*Calca*/CGRP^ cells. In the CeA, neurons projecting to the PB^*Calca*/CGRP^ neurons are mainly in the CeL whereas PB^*Calca*/CGRP^ terminals are concentrated in the central nucleus amygdala, capsular part (CeC). Future studies could establish the circuitry through which the PB^*Calca*/CGRP^ neurons interact with these extended amygdala neurons.

Still, not all of these connections are reciprocal. For example, rabies tracing shows that the NTS directly innervates the PB^*Calca*/CGRP^ neurons, but projections from the PB^*Calca*/CGRP^ neurons back to the NTS are rare (Huang et al., 2021b). In addition, the PB^*Calca*/CGRP^ neurons innervate many thalamic regions including somatosensory regions (Yasui et al., 1989; Bester et al., 1999; Krout and Loewy, 2000; Huang et al., 2021b), but we saw no evidence for thalamic projections back to the PB^*Calca*/CGRP^ neurons. Perhaps the PB^*Calca*/CGRP^ neurons interact closely with other limbic regions such as the extended amygdala, but for sensory signals, they may simply integrate and pass along information without much feedback.

### 4.2 Functional perspectives on inputs to the PB^*Calca*/CGRP^ neurons

As demonstrated previously, we found that the PB^*Calca*/CGRP^ cells are directly innervated by brain regions relaying noxious sensory stimuli (Palmiter, 2018). Conditional rabies tracing labeled many neurons in the NTS, and conditional anterograde tracing in *vGLUT2-Cre* mice showed that glutamatergic NTS nerve terminals closely appose PB^*Calca*/CGRP^ cells. The NTS inputs carry vagal signals related to nausea and conditioned food aversion (Huang et al., 2019; Qiu et al., 2023), and chemo- or photoactivation of the PB^*Calca*/CGRP^ neurons substantially reduces food intake (Wu et al., 2012; Carter et al., 2013). Furthermore, satiety signals from a full stomach increase the activity of PB^*Calca*/CGRP^ neurons via DBH and CCK neurons of the NTS (Roman et al., 2016). Conversely, inactivating the PB^*Calca*/CGRP^ neurons with tetanus toxin increases meal size, indicating their importance in spontaneous feeding as well (Roman et al., 2016). Aversion to certain food probably develops via projections to the extended amygdala as photostimulation of PB^*Calca*/CGRP^ axons in the BST and CeA produces taste aversion (Chen et al., 2018).

Similarly, the PB^*Calca*/CGRP^ neurons are essential for behavioral responses to pain. Conditional rabies tracing labeled many neurons in the trigeminal and reticular nuclei, and future work will likely show direct inputs from the dorsal horn and trigeminal ganglia as well (Rodriguez et al., 2017). Trigeminal pain is perceived more intensely than bodily pain (Rodriguez et al., 2017). With injections of CTb into the lateral PB, we find that the vast majority of CTb-labeled spinal trigeminal neurons are glutamatergic, suggesting they convey pain signals which excite the PB^*Calca*/CGRP^ neurons. Inputs from the medullary reticular formation may also relay pain signals to the PB^*Calca*/CGRP^ cells (Martins and Tavares, 2017). Notably, neuropathic pain from the sciatic nerve triggers brief arousals from sleep, and ablation of the PB^*Calca*/CGRP^ neurons prevents these awakenings (Alexandre, 2024).

These and many other studies show that the PB^*Calca*/CGRP^ neurons are essential for relaying signals related to noxious stimuli to the forebrain, but considering the variety of non-sensory, direct forebrain afferents to these cells, they probably also function within networks that shape behavioral responses. For example, signals from the extended amygdala and hypothalamus likely provide more complex information related to fear, hunger, and other internal states, and behavioral responses to fear illustrate how the PB^*Calca*/CGRP^ neurons can act independently of sensory stimuli. Mice often exhibit neophobia, and the presentation of a novel food initially increases PB^*Calca*/CGRP^ neuron activity, causing the mice to avoid the food, though this activity fades as the mice become acclimated to the food (Campos et al., 2016, 2018). Inactivation of the CGRP cells reduces this neophobia to novel food (Palmiter, 2018). In this case, the threat (novel food) is only perceived to be dangerous, suggesting that the signals driving this come from the forebrain rather than sensory afferents. Further supporting this idea, foot shock can activate the PB^*Calca*/CGRP^ neurons, and after pairing a tone with a foot shock, even the tone alone is enough to activate the PB^*Calca*/CGRP^ neurons (Sato et al., 2015; Palmiter, 2018). The fear-related signals likely come from the extended amygdala, and as the inputs from the BST and CeA are GABAergic, we propose that fear reduces activity in these amygdala neurons, thus disinhibiting the PB^*Calca*/CGRP^ neurons. Perhaps these GABAergic CeA neurons include the population expressing protein kinase C-delta and the oxytocin receptor which have anxiolytic effects (Haubensak et al., 2010). These forebrain fear circuits may engage the PB^*Calca*/CGRP^ neurons to drive fear and aversion behaviors, or perhaps the PB^*Calca*/CGRP^ neurons actually contribute to the emotional state itself.

### 4.3 Limitations

To map all inputs to the lateral PB, we injected a small volume of CTb, ensuring that the injection site was limited to this region. Though the injection site was primarily centered in the lateral PB, some mice showed slight extension into the adjacent medial PB and cerebellum. Still, the overall pattern is quite similar to that reported previously (Herbert et al., 1990; Moga et al., 1990; Tokita et al., 2009).

In addition, we used reporter mice in which the vGAT or vGLUT2 promoters drive Cre expression, resulting in GFP expression in these cells, yet Cre activity early in development permanently labels these cells, and their adult neurochemical phenotype may differ. For example, during development, many cortical neurons express vGLUT2 but as adults, they express vGLUT1 (Boulland et al., 2004b; Nakamura et al., 2005; Berry et al., 2012). In vGLUT2 reporter mice, we found that CTb labeled many GFP-containing cortical neurons, but as shown in Figure 7, these green cells actually express vGLUT1; the claim that these cells are glutamatergic still holds, but observations in reporter mice should be confirmed with additional methods.

We used conditional rabies virus tracing to map monosynaptic inputs to PB^*Calca*/CGRP^ neurons in the external lateral PB. While this method is intended to identify all primary input regions, it failed to label some known inputs to the PB^*Calca*/CGRP^ cells and labeled fewer neurons than expected. For instance, PB^*Calca*/CGRP^ neurons respond to norepinephrine and serotonin from the ventrolateral medulla, and raphe nuclei (Boccia et al., 2020; Kaur et al., 2020), yet we observed no rabies labeling in these or other monoaminergic nuclei. This limitation is likely inherent to the rabies method, as the transfer of rabies virus from a starter cell to afferent terminals may require direct, classical symmetric and asymmetric synapses, yet monoaminergic fibers often release neurotransmitters *en passant* with volume transmission (Toyoda et al., 2022). This general limitation of the rabies method is rarely mentioned but likely significant and probably applies to some peptidergic inputs as well. Moreover, the efficiency of labeling inputs depends on the levels of TVA and especially RG expression; if a starter cell expresses one or both helper proteins weakly, the number of rabies-labeled input cells can be low (Callaway and Luo, 2015).

It should also be noted that not all *Calca*-expressing neurons in the PB are CGRP-immunoreactive; neurons of the PBel contain substantial CGRP which can be easily detected with immunostaining, but *Calca*-expressing neurons of the medial PB and KF contain very little CGRP immunoreactivity. Recent MERFISH experiments suggest these cell groups differ in gene expression, but it remains unknown if these groups differ in their inputs or are functionally distinct. Indeed, Nardone and colleagues described *Calca* expression in two clusters at the level of the KF. These KF clusters show lower levels of *Calca* transcript compared to the PB *Calca*+/*Il20ra*+ cluster and are smaller in size. Among these clusters, *Onecut3* emerged as the most selective marker for these two KF *Calca* clusters, which are selectively marked by the expression of *Ebf2* and *Chst9*, allowing discrimination from the PB *Calca*+/*Il20ra*+ cluster (Pauli et al., 2022; Nardone et al., 2024).

## 5 Future Direction and Conclusions

Using a variety of techniques, we mapped the inputs to the PB^*Calca*/CGRP^ neurons, which are essential for waking in response to hypercarbia, suppressing appetite, and forming conditioned taste aversions. A key goal of future studies will be to use the results of RNAseq studies to intersectionally target these subpopulations of PB^*Calca*/CGRP^ neurons to determine if they are anatomically and functionally distinct. Of interest are the subgroup of external lateral PB^*Calca*/CGRP^ neurons that express the norepinephrine transporter, *Slc6a2*. These neurons are located in the dorsal part of the external lateral PB^*Calca*/CGRP^ neurons. Our CRACM studies indicate that CeA GABA neurons primarily project to and inhibit the external lateral PB^*Calca*/CGRP^ neurons dorsally located. In addition, it will be important to map inputs from the spinal cord and monitor the activity of key inputs using photometry and similar methods to begin modeling how these inputs influence the activity of the PB^*Calca*/CGRP^ neurons in response to stimuli such as hypercarbia, pain and cold, and across behavioral states. Optogenetic and chemogenetic studies on the terminals of these projecting neurons could further shed light on the roles of afferent projections under different aversive conditions in which PB^*Calca*/CGRP^ neurons are involved. These approaches should improve our understanding of the functions of this key neuronal population.

## Acknowledgments

We thank Sathyajit Bandaru for technical assistance with in situ hybridization and other histological procedures, and Clifford Saper for thoughtful feedback on this study.

## Authors Contribution

TES and MK planned the experiments; MK performed stereotaxic injections; MK and BF performed histologic staining and slide-scanning microscopy; RDL and EA conducted the CRACM experiments, analyzed the data, and drafted the CRACM figures; MK and TES analyzed data and wrote the manuscript; RDL and EA contributed to and edited the manuscript. MK drafted and edited all figures with TES; TES supervised the project. All authors reviewed and discussed the results and contributed to the final manuscript.

## REFERENCES

Alexandre C, Miracca G, Holanda VD, Sharma A, Kourbanova K, Ferreira A, Bicca MA, Zeng X, Nassar VA, Lee S, Kaur S, Sarma SV, Sacré P, Scammell TE, Woolf CJ, Latremoliere A. 2024. Nociceptor spontaneous activity is responsible for fragmenting non–rapid eye movement sleep in mouse models of neuropathic pain. Sci Transl Med 16:eadg3036.

Anaclet C, De Luca R, Venner A, Malyshevskaya O, Lazarus M, Arrigoni E, Fuller PM. 2018. Genetic Activation, Inactivation, and Deletion Reveal a Limited And Nuanced Role for Somatostatin-Containing Basal Forebrain Neurons in Behavioral State Control. J Neurosci 38:5168–5181.

Barik A, Sathyamurthy A, Thompson J, Seltzer M, Levine A, Chesler A. 2021. A spinoparabrachial circuit defined by Tacr1 expression drives pain. eLife 10:e61135.

Berry CT, Sceniak MP, Zhou L, Sabo SL. 2012. Developmental Up-Regulation of Vesicular Glutamate Transporter-1 Promotes Neocortical Presynaptic Terminal Development. PLoS ONE 7:e50911.

Bester H, Bourgeais L, Villanueva L, Besson JM, Bernard JF. 1999. Differential projections to the intralaminar and gustatory thalamus from the parabrachial area: a PHA-L study in the rat. J Comp Neurol 405:421–449.

Boccia L, Le Foll C, Lutz TA. 2020. Noradrenaline signaling in the LPBN mediates amylin’s and salmon calcitonin’s hypophagic effect in male rats. FASEB J 34:15448–15461.

Bochorishvili G, Stornetta RL, Coates MB, Guyenet PG. 2012. Pre-Bötzinger Complex Receives Glutamatergic Innervation From Galaninergic and Other Retrotrapezoid Nucleus Neurons. J Comp Neurol 520:1047–1061.

Boulland J-L, Qureshi T, Seal RP, Rafiki A, Gundersen V, Bergersen LH, Fremeau RT, Edwards RH, Storm-Mathisen J, Chaudhry FA. 2004a. Expression of the vesicular glutamate transporters during development indicates the widespread corelease of multiple neurotransmitters. J Comp Neurol 480:264–280.

Boulland J-L, Qureshi T, Seal RP, Rafiki A, Gundersen V, Bergersen LH, Fremeau RT, Edwards RH, Storm-Mathisen J, Chaudhry FA. 2004b. Expression of the vesicular glutamate transporters during development indicates the widespread corelease of multiple neurotransmitters. J Comp Neurol 480:264–280.

Callaway EM, Luo L. 2015. Monosynaptic Circuit Tracing with Glycoprotein-Deleted Rabies Viruses. J Neurosci Off J Soc Neurosci 35:8979–8985.

Campos CA, Bowen AJ, Han S, Wisse BE, Palmiter RD, Schwartz MW. 2017. Cancer-induced anorexia and malaise are mediated by CGRP neurons in the parabrachial nucleus. Nat Neurosci 20:934–942.

Campos CA, Bowen AJ, Roman CW, Palmiter RD. 2018. Encoding of danger by parabrachial CGRP neurons. Nature 555:617–622.

Campos CA, Bowen AJ, Schwartz MW, Palmiter RD. 2016. Parabrachial CGRP Neurons Control Meal Termination. Cell Metab 23:811–820.

Carter ME, Han S, Palmiter RD. 2015. Parabrachial calcitonin gene-related peptide neurons mediate conditioned taste aversion. J Neurosci Off J Soc Neurosci 35:4582–4586.

Carter ME, Soden ME, Zweifel LS, Palmiter RD. 2013. Genetic identification of a neural circuit that suppresses appetite. Nature 503:111–114.

Chen JY, Campos CA, Jarvie BC, Palmiter RD. 2018. Parabrachial CGRP Neurons Establish and Sustain Aversive Taste Memories. Neuron 100:891–899.e5.

De Luca R, Nardone S, Grace KP, Venner A, Cristofolini M, Bandaru SS, Sohn LT, Kong D, Mochizuki T, Viberti B, Zhu L, Zito A, Scammell TE, Saper CB, Lowell BB, Fuller PM, Arrigoni E. 2022. Orexin neurons inhibit sleep to promote arousal. Nat Commun 13:4163.

Feil K, Herbert H. 1995. Topographic organization of spinal and trigeminal somatosensory pathways to the rat parabrachial and Kölliker-Fuse nuclei. J Comp Neurol 353:506–528.

Geerling JC, Kim M, Mahoney CE, Abbott SBG, Agostinelli LJ, Garfield AS, Krashes MJ, Lowell BB, Scammell TE. 2016. Genetic identity of thermosensory relay neurons in the lateral parabrachial nucleus. Am J Physiol - Regul Integr Comp Physiol 310:R41–R54.

Grady F, Peltekian L, Iverson G, Geerling JC. 2020. Direct Parabrachial-Cortical Connectivity. Cereb Cortex N Y N 1991 30:4811–4833.

Han S, Soleiman MT, Soden ME, Zweifel LS, Palmiter RD. 2015. Elucidating an Affective Pain Circuit that Creates a Threat Memory. Cell 162:363–374.

Haubensak W, Kunwar PS, Cai H, Ciocchi S, Wall NR, Ponnusamy R, Biag J, Dong H-W, Deisseroth K, Callaway EM, Fanselow MS, Lüthi A, Anderson DJ. 2010. Genetic dissection of an amygdala microcircuit that gates conditioned fear. Nature 468:270–276.

Herbert H, Moga MM, Saper CB. 1990. Connections of the parabrachial nucleus with the nucleus of the solitary tract and the medullary reticular formation in the rat. J Comp Neurol 293:540–580.

Huang D, Grady FS, Peltekian L, Geerling JC. 2021a. Efferent projections of Vglut2, Foxp2, and Pdyn parabrachial neurons in mice. J Comp Neurol 529:657– 693.

Huang D, Grady FS, Peltekian L, Laing JJ, Geerling JC. 2021b. Efferent Projections of CGRP/Calca-expressing Parabrachial Neurons in Mice. J Comp Neurol 529:2911–2957.

Huang T, Lin S-H, Malewicz NM, Zhang Y, Zhang Y, Goulding M, LaMotte RH, Ma Q. 2019. Identifying the pathways required for coping behaviours associated with sustained pain. Nature 565:86–90.

Hull C, Isaacson JS, Scanziani M. 2009. Postsynaptic mechanisms govern the differential excitation of cortical neurons by thalamic inputs. J Neurosci Off J Soc Neurosci 29:9127–9136.

Hurley KM, Herbert H, Moga MM, Saper CB. 1991. Efferent projections of the infralimbic cortex of the rat. J Comp Neurol 308:249–276.

Jo A, Xu J, Deniz S, Cherian S, DeVries SH, Zhu Y. 2018. Intersectional Strategies for Targeting Amacrine and Ganglion Cell Types in the Mouse Retina. Front Neural Circuits 12:66.

Kang SJ, Liu S, Ye M, Kim D-I, Pao GM, Copits BA, Roberts BZ, Lee K-F, Bruchas MR, Han S. 2022. A central alarm system that gates multi-sensory innate threat cues to the amygdala. Cell Rep 40:111222.

Kaur S, De Luca R, Khanday MA, Bandaru SS, Thomas RC, Broadhurst RY, Venner A, Todd WD, Fuller PM, Arrigoni E, Saper CB. 2020. Role of serotonergic dorsal raphe neurons in hypercapnia-induced arousals. Nat Commun 11:2769.

Kaur S, Nicole L, Sela Y, Lima J, Thomas R, Bandaru S, Saper C. 2023. Lateral parabrachial FoxP2 neurons regulate respiratory responses to hypercapnia. Res Sq:rs.3.rs-2865756.

Kaur S, Pedersen NP, Yokota S, Hur EE, Fuller PM, Lazarus M, Chamberlin NL, Saper CB. 2013. Glutamatergic signaling from the parabrachial nucleus plays a critical role in hypercapnic arousal. J Neurosci Off J Soc Neurosci 33:7627–7640.

Kaur S, Saper C. 2021. FoxP2 cells in the lateral parabrachial area may drive respiratory responses to hypercapnia. FASEB J [Internet] 35. Available from: https://onlinelibrary.wiley.com/doi/abs/10.1096/fasebj.2021.35.S1.02023

Kaur S, Wang JL, Ferrari L, Thankachan S, Kroeger D, Venner A, Lazarus M, Wellman A, Arrigoni E, Fuller PM, Saper CB. 2017. A Genetically Defined Circuit for Arousal from Sleep during Hypercapnia. Neuron 96:1153-1167.e5.

Kirouac GJ, Li S, Li S. 2022. Convergence of monosynaptic inputs from neurons in the brainstem and forebrain on parabrachial neurons that project to the paraventricular nucleus of the thalamus. Brain Struct Funct 227:2409–2437.

Krashes MJ, Shah BP, Madara JC, Olson DP, Strochlic DE, Garfield AS, Vong L, Pei H, Watabe-Uchida M, Uchida N, Liberles SD, Lowell BB. 2014. An excitatory paraventricular nucleus to AgRP neuron circuit that drives hunger. Nature 507:238–242.

Krout KE, Loewy AD. 2000. Parabrachial nucleus projections to midline and intralaminar thalamic nuclei of the rat. J Comp Neurol 428:475–494.

Lee S, Augustine V, Zhao Y, Ebisu H, Ho B, Kong D, Oka Y. 2019. Chemosensory modulation of neural circuits for sodium appetite. Nature 568:93–97.

Lynch N, Thomas R, Saper C, Kaur S. 2022. 0171 Parabrachial-CGRP neurons regulate awakenings to pain stimulus. Sleep 45:A79.

Martins I, Tavares I. 2017. Reticular Formation and Pain: The Past and the Future. Front Neuroanat 11:51.

McDonald AJ. 1982. Cytoarchitecture of the central amygdaloid nucleus of the rat. J Comp Neurol 208:401– 418.

McDonald AJ. 1983. Neurons of the bed nucleus of the stria terminalis: a golgi study in the rat. Brain Res Bull 10:111–120.

Miller RL, Knuepfer MM, Wang MH, Denny GO, Gray PA, Loewy AD. 2012. Fos-activation of FoxP2 and Lmx1b neurons in the parabrachial nucleus evoked by hypotension and hypertension in conscious rats. Neuroscience 218:110–125.

Moga MM, Herbert H, Hurley KM, Yasui Y, Gray TS, Saper CB. 1990. Organization of cortical, basal forebrain, and hypothalamic afferents to the parabrachial nucleus in the rat. J Comp Neurol 295:624– 661.

Nakamura K, Hioki H, Fujiyama F, Kaneko T. 2005. Postnatal changes of vesicular glutamate transporter (VGluT)1 and VGluT2 immunoreactivities and their colocalization in the mouse forebrain. J Comp Neurol 492:263–288.

Nardone S, De Luca R, Zito A, Klymko N, Nicoloutsopoulos D, Amsalem O, Brannigan C, Resch JM, Jacobs CL, Pant D, Veregge M, Srinivasan H, Grippo RM, Yang Z, Zeidel ML, Andermann ML, Harris KD, Tsai LT, Arrigoni E, Verstegen AMJ, Saper CB, Lowell BB. 2024. A spatially-resolved transcriptional atlas of the murine dorsal pons at single-cell resolution. Nat Commun 15:1966.

Norgren R, Leonard CM. 1971. Taste pathways in rat brainstem. Science 173:1136–1139.

Palmiter RD. 2018. The parabrachial nucleus: CGRP neurons function as a general alarm. Trends Neurosci 41:280–293.

Pauli JL, Chen JY, Basiri ML, Park S, Carter ME, Sanz E, McKnight GS, Stuber GD, Palmiter RD. 2022. Molecular and anatomical characterization of parabrachial neurons and their axonal projections. eLife 11:e81868.

Paxinos G. 2004. The Rat Nervous System. Elsevier Academic Press.

Paxinos G, Franklin KBJ. 2004. The Mouse Brain in Stereotaxic Coordinates. Gulf Professional Publishing.

Qiu W, Hutch CR, Wang Y, Wloszek J, Rucker RA, Myers MG, Sandoval D. 2023. Multiple NTS neuron populations cumulatively suppress food intake. eLife 12:e85640.

Roberts GW, Woodhams PL, Polak JM, Crow TJ. 1982. Distribution of neuropeptides in the limbic system of the rat: the amygdaloid complex. Neuroscience 7:99–131.

Rodriguez E, Sakurai K, Xu J, Chen Y, Toda K, Zhao S, Han B-X, Ryu D, Yin H, Liedtke W, Wang F. 2017. A craniofacial-specific monosynaptic circuit enables heightened affective pain. Nat Neurosci 20:1734–1743.

Roman CW, Derkach VA, Palmiter RD. 2016. Genetically and functionally defined NTS to PBN brain circuits mediating anorexia. Nat Commun 7:11905.

Saper CB. 1982. Reciprocal parabrachial-cortical connections in the rat. Brain Res 242:33–40.

Saper CB. 2016. The House Alarm. Cell Metab 23:754– 755.

Sato M, Ito M, Nagase M, Sugimura YK, Takahashi Y, Watabe AM, Kato F. 2015. The lateral parabrachial nucleus is actively involved in the acquisition of fear memory in mice. Mol Brain 8:22.

Schwaber JS, Kapp BS, Higgins GA, Rapp PR. 1982. Amygdaloid and basal forebrain direct connections with the nucleus of the solitary tract and the dorsal motor nucleus. J Neurosci Off J Soc Neurosci 2:1424–1438.

Shin J-W, Geerling JC, Loewy AD. 2008. Inputs to the ventrolateral bed nucleus of the stria terminalis. J Comp Neurol 511:628–657.

Simon MM, Greenaway S, White JK, Fuchs H, Gailus-Durner V, Wells S, Sorg T, Wong K, Bedu E, Cartwright EJ, Dacquin R, Djebali S, Estabel J, Graw J, Ingham NJ, Jackson IJ, Lengeling A, Mandillo S, Marvel J, Meziane H, Preitner F, Puk O, Roux M, Adams DJ, Atkins S, Ayadi A, Becker L, Blake A, Brooker D, Cater H, Champy M-F, Combe R, Danecek P, di Fenza A, Gates H, Gerdin A-K, Golini E, Hancock JM, Hans W, Hölter SM, Hough T, Jurdic P, Keane TM, Morgan H, Müller W, Neff F, Nicholson G, Pasche B, Roberson L-A, Rozman J, Sanderson M, Santos L, Selloum M, Shannon C, Southwell A, Tocchini-Valentini GP, Vancollie VE, Westerberg H, Wurst W, Zi M, Yalcin B, Ramirez-Solis R, Steel KP, Mallon A-M, de Angelis MH, Herault Y, Brown SDM. 2013. A comparative phenotypic and genomic analysis of C57BL/6J and C57BL/6N mouse strains. Genome Biol 14:R82.

Swanson LW. 2004. Brain Maps: Structure of the Rat Brain. Gulf Professional Publishing.

Tokita K, Inoue T, Boughter JD. 2009. Afferent connections of the parabrachial nucleus in C57BL/6J mice. Neuroscience 161:475–488.

Toyoda H, Won J, Kim W, Kim H, Davy O, Saito M, Kim D, Tanaka T, Kang Y, Oh SB. 2022. The Nature of Noradrenergic Volume Transmission From Locus Coeruleus to Brainstem Mesencephalic Trigeminal Sensory Neurons. Front Cell Neurosci [Internet] 16. Available from: https://www.frontiersin.org/articles/10.3389/fncel.2022.841239

Verstegen AMJ, Vanderhorst V, Gray PA, Zeidel ML, Geerling JC. 2017. Barrington’s nucleus: Neuroanatomic landscape of the mouse “pontine micturition center.” J Comp Neurol 525:2287–2309.

Vong L, Ye C, Yang Z, Choi B, Chua S, Lowell BB. 2011. Leptin action on GABAergic neurons prevents obesity and reduces inhibitory tone to POMC neurons. Neuron 71:142–154.

Woodhams PL, Roberts GW, Polak JM, Crow TJ. 1983. Distribution of neuropeptides in the limbic system of the rat: the bed nucleus of the stria terminalis, septum and preoptic area. Neuroscience 8:677–703.

Wu Q, Clark MS, Palmiter RD. 2012. Deciphering a neuronal circuit that mediates appetite. Nature 483:594– 597.

Yasui Y, Saper CB, Cechetto DF. 1989. Calcitonin gene-related peptide immunoreactivity in the visceral sensory cortex, thalamus, and related pathways in the rat. J Comp Neurol 290:487–501.

